# Increased functional integration of emotional control network across the adult lifespan

**DOI:** 10.1101/2024.04.10.588823

**Authors:** Leona Rahel Bätz, Shuer Ye, Melissa Schubbert, Xiaqing Lan, Maryam Ziaei

## Abstract

Emotional wellbeing has often been shown to improve across the adult lifespan. The corresponding effects, however, in the emotion regulation brain networks remain underexplored. By utilizing large-scale datasets such as the Human Connectome Project (HCP-Aging, N=621, 349 females) and Cambridge Centre for Ageing and Neuroscience (Cam-CAN, N=333, 155 females), we were able to investigate how emotion regulation networks’ functional topography differs across the entire adult lifespan. Based on previous meta-analytic work that identified four large-scale functional brain networks involved in emotion generation and regulation, we investigated the association between the integration of these emotion regulation networks and measures of mental wellbeing with age in the HCP-Aging dataset. We found an increased functional integration of the emotional control network among older adults, which was replicated using the Cam-CAN dataset. Further we found that the interoceptive network, which is mediating emotion generative and regulative processes and carries our introspective and reflective functions, is less integrated in higher age. Our study highlights the importance of identifying topological changes in the functional emotion network architecture across the lifespan, as it allows for a better understanding of functional brain network changes that accompany emotional aging.

**Highlights:**

- We aimed to identify age-related differences in the functional integration of large-scale emotion regulation brain networks across the adult lifespan, and its implications for well-being.
- Frontal emotion control network showed increased while interoception network showed decreased functional integration with higher age in the HCP-Aging dataset.
- Increased integration of emotion control network was replicated and validated in the Cam-CAN dataset.

## 1 Introduction

Growing old is often accompanied by a range of cognitive, physical and behavioral changes (Spreng & Turner, 2019). The perception and regulation of emotion are also subject to these changes in late adulthood (Charles, 2010; Mather, 2012, 2024). However, while cognitive capabilities tend to decline, this is not necessarily true for emotional well-being (Charles et al., 2009; Livingstone & Isaacowitz, 2021; Reed & Carstensen, 2012; Reuter-Lorenz & Lustig, 2005; Wolfe & Isaacowitz, 2022). Longitudinal studies have revealed that aging is associated with overall improved emotional well-being and greater emotional stability (Carstensen et al., 2011). It is debated whether this can or cannot be attributed to better emotion regulation (Urry & Gross, 2010). However, older adults have been shown to select situations that avoid conflict or negative affect (Birditt et al., 2005; Charles et al., 2009; Isaacowitz & English, 2024), deploy more attention to positive emotions (Isaacowitz et al., 2006), and prioritize effective emotion regulation more than their younger counterparts (Arioli et al., 2018; English & Carstensen, 2014; Lawton et al., 1992; Nolen-Hoeksema & Aldao, 2011; Ziaei et al., 2017). While the mechanisms underlying such age-related shift is not completely understood, older adults maintain an absence of negative affect more consistently and can recover more quickly from negative emotional states compared to younger adults (Carstensen et al., 2000; Hay & Diehl, 2011; Isaacowitz & English, 2024). The Socioemotional Selectivity Theory explains this bias by stating that older adults shift their focus from future-oriented goals to present-oriented ones, with positive stimuli holding a more immediate appeal (Carstensen et al., 1999). The physiological hypothesis of emotional aging (MacCormack et al., 2020; 2021; 2023) proposes that age-related changes in bodily and neural physiology—particularly in interoceptive processing—contribute to differences in emotional experience and regulation across adulthood. This model suggests that alterations in afferent bodily signals and interoceptive representation shape emotional awareness and reactivity, complementing the motivational focus described by SST. While the selective engagement hypothesis postulates that due to older adults’ limited cognitive capabilities compared to younger adults, they are more astute in expending these finite resources (Hess, 2014). Hence, they allocate their resources more discerningly, leading to better performance on tasks with personal relevance, significance, or social implications (Carstensen et al., 2011).

Accompanied by these motivational changes and their reflection in behavior, several changes are occurring at the neural level (MacCormack et al., 2020; Mather, 2016). Investigating such changes in the brain will help to better understand the motivational and behavioral changes observed across the lifespan. For instance, older adults exhibit increased activity of the prefrontal cortex (PFC) and reduced activity of the amygdala in comparison to younger adults in several tasks that require emotional processing (Mather, 2012; Nashiro et al., 2012; Ziaei et al., 2017), reflecting the potential inhibitory or regulatory control from the PFC to the amygdala. Emotion regulation is a complex process, and while some of the improved emotion regulation in later life was attributed to the increased recruitment of frontal areas during this process (Pessoa, 2008; St. Jacques et al., 2010; Tomasi & Volkow, 2012), the impact of this observed recruitment of frontal areas on the rest of the brain has yet to be fully understood. The complex interplay between major nodes in the affect system, including the PFC and amygdala, insula, anterior cingulate cortex, and ventral striatum is critical for better understanding, experiencing, and regulating emotions (Ahmed et al., 2015; Ghashghaei et al., 2007). When investigating the emotional response specifically on a network level, a subsystem of four brain networks that are specifically involved in emotion generation and regulation has been identified in a comprehensive meta-analytic study, known as meta-analytic groupings (MAGs) (Morawetz et al., 2020). To identify the MAGs, *K*-means clustering was performed on imaging data from 385 fMRI experiments from 107 published papers that investigated emotion generative processes vs. emotion regulatory processes, with forward inference to establish functional fingerprints of the MAGs. In total, four networks were identified, referenced as MAG1 to MAG4, which have different overall roles in emotional responses. MAG1 is involved in emotional control and consists of more frontal regions. MAG2 is also a mostly frontal and left lateralized network, which is involved in emotion regulation and verbalization. MAG3 involves mostly subcortical areas and is associated with emotion regulation. MAG4 comprises both cortical and subcortical regions, like the thalamus and insula, and is considered to be involved in the detection of emotional stimuli and interoception (Morawetz et al., 2020). The first and second MAG (i.e., MAG1 and MAG2) comprise mostly cortical areas including parts of the PFC, while the third and fourth MAG (i.e., MAG3 and MAG4) also include subcortical areas like the bilateral amygdala. Morawetz et al identified the functional fingerprints of the MAGs describing that MAG1’s role is predominantly in working memory, response inhibition, and reasoning; thus, it can be referred to as exhibiting emotional control functions. MAG2 also shows a strong association with cognitive control, but it places a stronger focus on language compared to MAG1. On the other hand, MAG3 specializes in emotional reactivity and memory, while MAG4 has a more mixed functional fingerprint, being involved in both emotion generative and regulatory processes. MAG4’s activation is associated with pain perception, interoception, and perception of the emotional stimulus in both generative and regulatory phases. Thus, MAG3 can be simplified to be an emotion generation network, while MAG4 can be defined as an interoception network.

Generally, besides the MAGs, other large-scale brain networks are not exempt from age-related changes, as overall higher network integration in these networks in later life stage has been reported (Chan et al., 2014; Ferreira & Busatto, 2013; Grady et al., 2016; L. He et al., 2020; Onoda & Yamaguchi, 2013). High functional integration of a brain network describes that it is strongly connected across the whole cortex while low functional integration describes high modularity of the network and richer within connections (Sporns, 2013). Increased integration of the frontoparietal control network (FPN), default mode network (DMN), and dorsal attention network (DAN) predicts poorer working memory and decreased processing speed in late adulthood (Ng et al., 2016; Salami et al., 2018). However, higher integration may not only result in deficits regarding behavioral performance. Previous studies have shown that the age-related increases in integration of the salience network within the rest of the cortex (Voss et al., 2013), and enhanced integration between the executive control network and the rest of the cortex have been associated with improved life satisfaction in older adults (Lyoo & Yoon, 2017). Increased integration of specifically right hemispheric networks was further associated with greater working memory capacity in older adults, but not in younger (Crowell et al., 2020). Indicating that cognition in the aged brain might, to some degree, depends on or benefits from the network integration. However, increased integration at the network level might also be detrimental to the system, as it implies that in which more neural systems are activated in parallel reflecting a more diffuse, and nonspecific recruitment of brain systems. Other findings indicate as a consequence of an increased global integration in aging, older adults show slower cognitive processing (Imms et al., 2021; Xin Li et al., 2023). Generally, it can be inferred that changes at the network level can influence or even predict observable behavioral changes, although with differential impact depending on the emotional or cognitive nature of the functions (Chen et al., 2022; Levakov et al., 2021; Wei Li et al., 2023; Yu Li et al., 2022).

Alteration occurring within the functional topography of emotion regulation networks across the lifespan could therefore provide valuable insights into this nuanced picture of age-related shift in emotional processes and how it might be linked with emotional wellbeing in this population (Gurera & Isaacowitz, 2019; Isaacowitz & English, 2024; Thomas et al., 2016). However, the connections between emotion regulation network functional integration and mental wellbeing across the aging lifespan is yet to be fully examined. Therefore, the present study aims to investigate whether and how the emotion regulation networks’ topography changes across the adult lifespan, and whether the changes in the networks’ organization are associated with psychological outcomes. Given that there is a high correspondence between task and resting-state functional connectivity (Cole et al., 2016; Killgore et al., 2017; Krylova et al., 2021) we have focused on the topological characteristics of MAGs during resting state. By linking network connectivity of MAGs measured during rest to subsequent performance and regulation tendency outside the scanner, a recent study supports the idea that intrinsic functional architecture can serve as a trait-based predictor of emotional regulation outcomes (Morawetz et al., 2025). Previous research has linked functional connectivity in the MAGs to remission in major depressive disorder (Hang Wu et al., 2022), demonstrating that the MAGs themselves exhibit functional relevance for emotional and wellbeing-related domains, making them a suitable network model for the current study. Thus, we investigated the functional integration, using multiple metrices of functional integration, of the four large-scale networks (MAGs) across the adult lifespan with the available dataset of the Human Connectome Project - Aging (HCP-Aging), and the Cambridge Centre for Ageing and Neuroscience (Cam-CAN) dataset.

Considering the increased involvement of frontal areas during emotional processing and emotion regulation, specifically (J. U. Kim et al., 2019; Nashiro et al., 2012; Pessoa, 2008), we speculated that the predominantly frontal emotional control networks (such as MAGs 1 and 2) would become more integrated in late adulthood. The increased integration was expected to facilitate the inhibitory control required for emotion regulation, primarily exhibited by the frontal areas (Aron, 2007). Thus, we anticipated that frontal nodes of emotion regulation networks become more integrated with increasing age. Secondly, we hypothesized that differences in functional network integration would mediate age-related effects on perceived negative affects measured by psychological questionnaires. This was based on the idea that higher ability in regulating negative emotions could result in higher emotional well-being and lower stress (Extremera et al., 2020; Gross et al., 1997; Kraiss et al., 2020). Specifically, improved measures of emotional wellbeing (depression, anxiety, loneliness, stress) are mediated by network and nodal integration. Together, these analyses will provide additional insight into how functional emotion networks differ across the adult lifespan with specific details on the differences of functional integration, and association with psychological measures.

## 2 Material and methods

### 2.1 Participants

The data used for this study were obtained from the HCP-Aging (Bookheimer et al., 2019) and the Cam-CAN datasets (Shafto et al., 2014; Taylor et al., 2016). We utilized minimally processed imaging data and demographic information, which included the age, gender and years of education of each subject. Additionally, questionnaire data, such as stress scale, loneliness scale, and depression and anxiety scales from the NIH Toolbox (Hodes et al., 2013), were used to assess brain-behavior relationship (**Table 1**).

**Table 1:**
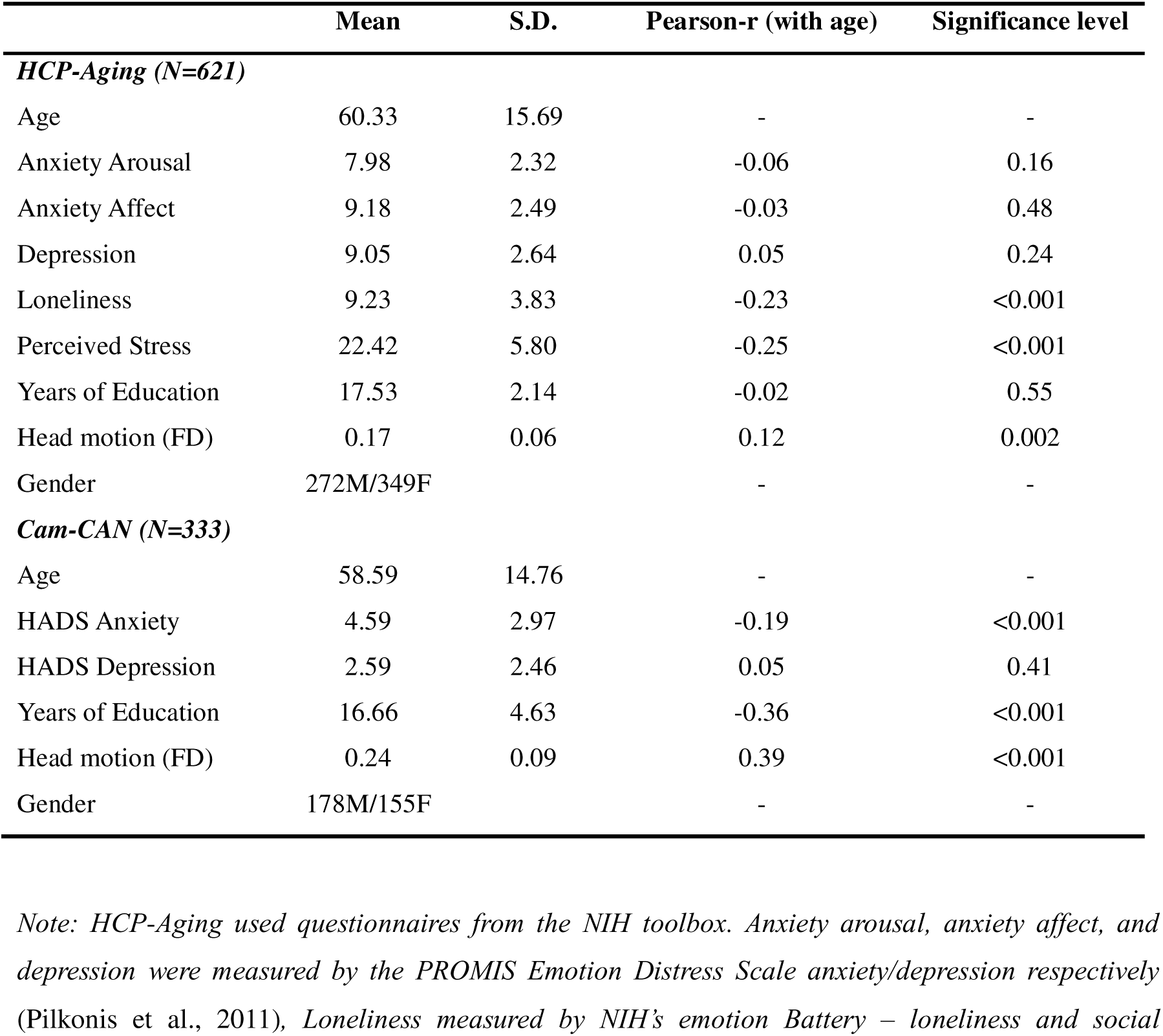
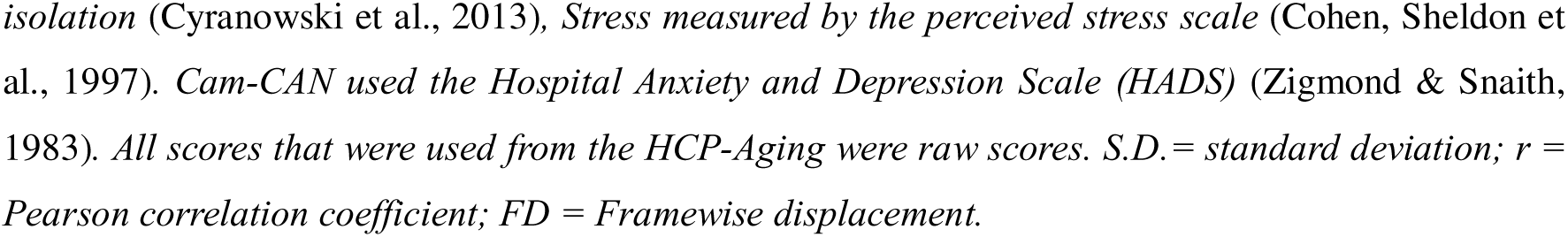
Descriptive Data from HCP-Aging and Cam-CAN.

|  | Mean | S.D. | Pearson-r (with age) | Significance level |
| --- | --- | --- | --- | --- |
| ***HCP-Aging (N=621)*** |  |  |  |  |
| Age | 60.33 | 15.69 | - | - |
| Anxiety Arousal | 7.98 | 2.32 | -0.06 | 0.16 |
| Anxiety Affect | 9.18 | 2.49 | -0.03 | 0.48 |
| Depression | 9.05 | 2.64 | 0.05 | 0.24 |
| Loneliness | 9.23 | 3.83 | -0.23 | <0.001 |
| Perceived Stress | 22.42 | 5.80 | -0.25 | <0.001 |
| Years of Education | 17.53 | 2.14 | -0.02 | 0.55 |
| Head motion (FD) | 0.17 | 0.06 | 0.12 | 0.002 |
| Gender | 272M/349F |  | - | - |
| ***Cam-CAN (N=333)*** |  |  |  |  |
| Age | 58.59 | 14.76 | - | - |
| HADS Anxiety | 4.59 | 2.97 | -0.19 | <0.001 |
| HADS Depression | 2.59 | 2.46 | 0.05 | 0.41 |
| Years of Education | 16.66 | 4.63 | -0.36 | <0.001 |
| Head motion (FD) | 0.24 | 0.09 | 0.39 | <0.001 |
| Gender | 178M/155F |  | - | - |

*Note: HCP-Aging used questionnaires from the NIH toolbox. Anxiety arousal, anxiety affect, and depression were measured by the PROMIS Emotion Distress Scale anxiety/depression respectively (Pilkonis et al., 2011), Loneliness measured by NIH's emotion Battery – loneliness and social*
*isolation* (Cyranowski et al., 2013), *Stress measured by the perceived stress scale* (Cohen, Sheldon et al., 1997). *Cam-CAN used the Hospital Anxiety and Depression Scale (HADS)* (Zigmond & Snaith, 1983). All scores that were used from the HCP-Aging were raw scores. *S.D.* = standard deviation; *r* = Pearson correlation coefficient; *FD* = Framewise displacement.

In total, after excluding participants with translation that exceeded 2 mm or rotation that exceeded 2° within a single TR, and structural abnormalities in the brain, 621 healthy participants (349 females) between the ages of 36 and 90 years old (Mean age = 60.3 ± 15.7) were included in the analyses. Subjects with suspected Alzheimer’s disease and neurological disorders (e.g., brain tumors, Parkinson’s disease, stroke) were excluded in both datasets. Additionally, all participants were drawn from the Human Connectome Project in Aging (HCP-A) and met the inclusion criteria established by the consortium. Subjects with suspected Alzheimer’s disease, mild cognitive impairment, or neurological disorders (e.g., stroke, Parkinson’s disease, brain tumors) were excluded. Additionally, participants diagnosed or treated for major psychiatric disorders (e.g., schizophrenia, bipolar disorder) or major depression lasting 12 months or more within the past five years were excluded. Cognitive status in HCP-A was determined using age-adjusted Montreal Cognitive Assessment (MoCA) scores and health records; only individuals within the normal cognitive range were included. While cognitive data (e.g., MoCA, memory, and executive function tests) are available from HCP-A, we restricted our analysis to the healthy aging cohort as defined by the project (Bookheimer et al., 2019). It should be noted that behavioral, cognitive, biosample, and multimodal MRI data for the HCP-Aging cohort were collected within the same testing wave. Participants completed a comprehensive assessment visit lasting up to approximately eight hours, typically within a single day (occasionally spread over two consecutive days for tolerance). When two days were required, behavioral assessments were conducted on the first day and imaging on the following day. Thus, all wellbeing and neuroimaging data used in this study were acquired within the same testing period (Bookheimer et al., 2019).

All primary analyses were carried out on the HCP-Aging and for validation purposes, we subsequently performed the same analyses on data from 333 participants between 36-88 years old (155 Female, mean age = 58.6 ± 14.7; Figure S1) from the Cam-CAN dataset using the same pipelines as for the HCP-Aging dataset. Gender distribution of both datasets in each age range is now reported in Figure S1. All participants in both the HCP-Aging and Cam-CAN datasets provided informed consent and their participation was voluntary.

### 2.2 Acquisition and preprocessing of neuroimaging data

The brain images from HCP-Aging were acquired using a 32-channel head coil on a 3T Siemens Prisma System across multiple centers (Washington University, University of Minnesota, Massachusetts General Hospital, Harvard University, University of California Los Angeles, Oxford University). Acquisition protocols were unified across all centers. T1 structural images in 0.8 mm resolution were acquired using the multi-echo MPRAGE sequence (TR = 2500 ms; voxel size = 0.8×0.8×0.8 mm, TI = 1000 ms; TE = 1.8/3.6/5.4/7.2 ms; and flip angle= 8°). Acquisition of functional data was done with a 2D multiband gradient echo planar imaging (EPI) sequence (TR = 800 ms; TE = 37 ms; 72 axial slices; voxel size = 2.0 × 2.0 × 2.0 mm; flip angle = 52°; multiband factor = 8). All participants completed four sessions of 6 minutes and five seconds of eyes-open resting-state scans, total of 30.4 minutes of resting-state scan. Sessions were conducted either on the same or consecutive days (detail of scanning parameters for HCP-Aging and Cam-CAN can be found in **Table S1**).

We acquired the minimally processed data from HCP-Aging, which included spatial artifact/distortion removal using the field maps calculated based on the difference in distortion between the two-phase encoding directions, surface generation, cross-modal registration, and alignment to standard space (Glasser et al., 2013). CONN Toolbox version 19v was used to perform additional in-house preprocessing steps comprised of spatial smoothing with an isotropic Gaussian kernel of 4 mm full width at half maximum, bandpass filtering at 0.008-0.09 Hz, denoising including anatomical component-based noise correction procedure (aComCor), motion regression with 12 regressors (6 motion parameters and their first-order derivatives), scrubbing (removal of a variable number of noise components based on outlier scans that either showed framewise displacement larger than 0.9 or BOLD signal changes over 5 S.D.), and linear detrending (Nieto-Castanon, 2020).

The four meta-analytic groupings (MAGs) used in this study were defined by Morawetz et al. (2020) through meta-analytic clustering of co-activation patterns across more than 1000 task-based emotion studies. Thus, the MAGs represent functionally defined networks implicated in emotion generation and regulation, rather than connectivity-based networks derived directly from the present dataset. We examined resting-state functional connectivity within and between these predefined emotion-related MAGs to assess how their intrinsic integration patterns vary with age. Averaged time courses of the masked voxels for each of the 36 ROIs of the MAGs were extracted using masks provided by Morawetz et al (Table 2). The masks contain the exact voxels identified in the original paper that established the MAGs (Morawetz et al., 2020). We decided to use these masks due to the non-spherical nature of many of these ROIs, making the signal localization more precise.

**Table 2.** MNI coordinates of 36 regions of interest from all four meta-analytic groupings (MAGs)

| Serial number | Hemisphere | Region | Volume (voxels) | MNI Coordinates |  |  |
| --- | --- | --- | --- | --- | --- | --- |
|  |  |  |  | x | y | z |
| MAG1 - Emotional Control |  |  |  |  |  |  |
| 1 | L | Superior Frontal Gyrus | 11704 | 0 | 24 | 50 |
| 2 | R | Middle Frontal Gyrus | 11024 | 40 | 24 | 42 |
| 3 | R | Inferior Parietal Lobule | 9968 | 58 | -52 | 38 |
| 4 | L | Inferior Parietal Lobule | 6216 | -58 | -50 | 44 |
| 5 | L | Middle Frontal Gyrus | 4664 | -36 | 52 | -2 |
| 6 | L | Middle Frontal Gyrus | 4288 | -42 | 14 | 48 |
| 7 | R | Middle Frontal Gyrus | 2792 | 42 | 46 | -8 |
| 8 | R | Insula | 2000 | 36 | 16 | 6 |
| 9 | R | Cingulate Gyrus | 1336 | 2 | -22 | 30 |
| 10 | R | Precuneus | 944 | 10 | -64 | 36 |
| MAG2 - Emotional Control (language) |  |  |  |  |  |  |
| 11 | L | Inferior Frontal Gyrus | 19464 | -46 | 24 | -8 |
| 12 | L | Superior Frontal Gyrus | 16592 | -4 | 10 | 62 |
| 13 | R | Inferior Frontal Gyrus | 6856 | 50 | 28 | -8 |
| 14 | L | Superior Temporal Gyrus | 6704 | -46 | -52 | 28 |
| 15 | L | Middle Temporal Gyrus | 5024 | -54 | -34 | -2 |
| 16 | L | Middle Frontal Gyrus | 4568 | -44 | 6 | 50 |
| 17 | L | Superior Frontal Gyrus | 3080 | -30 | 48 | 26 |
| 18 | L | Caudate | 1960 | -16 | 10 | 12 |
| 19 | R | Tuber | 1640 | 36 | -60 | -30 |
| MAG3 - Emotion Generation |  |  |  |  |  |  |
| 20 | L | Amygdala | 8640 | -22 | -4 | -16 |
| 21 | R | Amygdala | 6512 | 24 | -4 | -18 |
| 22 | R | Fusiform Gyrus | 4776 | 40 | -46 | -18 |
| 23 | R | Thalamus | 3528 | 6 | -26 | 0 |
| 24 | L | Fusiform Gyrus | 1256 | -38 | -54 | -14 |
| 25 | L | Parahippocampal Gyrus | 1216 | -22 | -28 | -4 |
| 26 | B | Medial Frontal Gyrus | 1016 | 0 | 54 | -10 |
| 27 | L | Inferior Occipital Gyrus | 912 | -42 | -76 | -6 |
| MAG4 - Interoception |  |  |  |  |  |  |
| 28 | L | Postcentral Gyrus | 4160 | -58 | -22 | 32 |
| 29 | L | Insula | 3752 | -44 | -4 | 10 |
| 30 | L | Superior Parietal Lobule | 2240 | -28 | -52 | 56 |
| 31 | R | Postcentral Gyrus | 1763 | 62 | -22 | 30 |
| 32 | L | Cuneus | 1224 | -10 | -76 | 22 |
| 33 | L | Middle Occipital Gyrus | 1152 | -48 | -74 | 2 |
| 34 | R | Thalamus | 1024 | 10 | -26 | -4 |
| 35 | R | Precuneus | 832 | 28 | -60 | 38 |
| 36 | R | Posterior Cingulate | 832 | 16 | -56 | 16 |

*Note: L=left; R=right; B=bilateral; MNI = Montreal Neurological Institute.*

### 2.3 Topological analyses of integration of MAG networks

We used resting state data from both HCP-Aging and Cam-CAN datasets to test our hypotheses. Integration refers to the capacity of the network to become interconnected and exchange information (Sporns, 2022). The term integration will refer to the degree to which a module of the graph has connections to the whole network in our case one MAG with all the MAGs together. Integration therefore always refers to the connection to the network comprising all 36 areas of all four MAGs. A module that is highly integrated exhibits extensive connections and interactions with other modules of the graph, enabling communication and coordination across the brain network. Two measurements of integration are given by the participation coefficient and intra- and inter-modular connectivity (i.e., modular interaction, Jinhui Wang et al., 2010).

To address differences in integration of emotion regulation functional networks and their respective areas (Hypotheses 1 and 2) across the lifespan, we employed a graph theoretical approach. Four meta-analytical groupings (**Figure 1A**) were chosen with 36 regions of interest (ROI), and ROI-wise time series were extracted by averaging the signal across voxels within the ROI (coordinates of all ROIs are presented in **Table 2**). Time courses of each ROI were correlated using Pearson correlation and then the resulting correlation matrices were *r-*to*-z* transformed. These *z*-transformed values were saved in 36×36 functional connectivity (FC) matrices. Subsequently, we constructed graphs from these FC matrices, where each node represented an ROI, and the edges represented functional connections. We specifically selected the top 30% highest positive values from the FC matrices, which were then represented as undirected unweighted edges in the graph. While the choice of a 30% sparsity level was based on the argument of its biological plausibility (Dennis et al., 2012; Sporns & Betzel, 2016), to ensure robustness of our results, we tested different sparsity values of 0.2, 0.25, 0.3 and 0.35 and our results remained consistent with what is reported with sparsity of 0.3. Each of the MAGs represented a functional module within the graph. In addition, while the size of ROIs was different, correlating the mean time-courses of all voxels in each ROI to construct FC matrices ensures that there is no bias in the signal from each ROI affecting the results (For workflow see **Figure 1B**).

**Figure 1.**
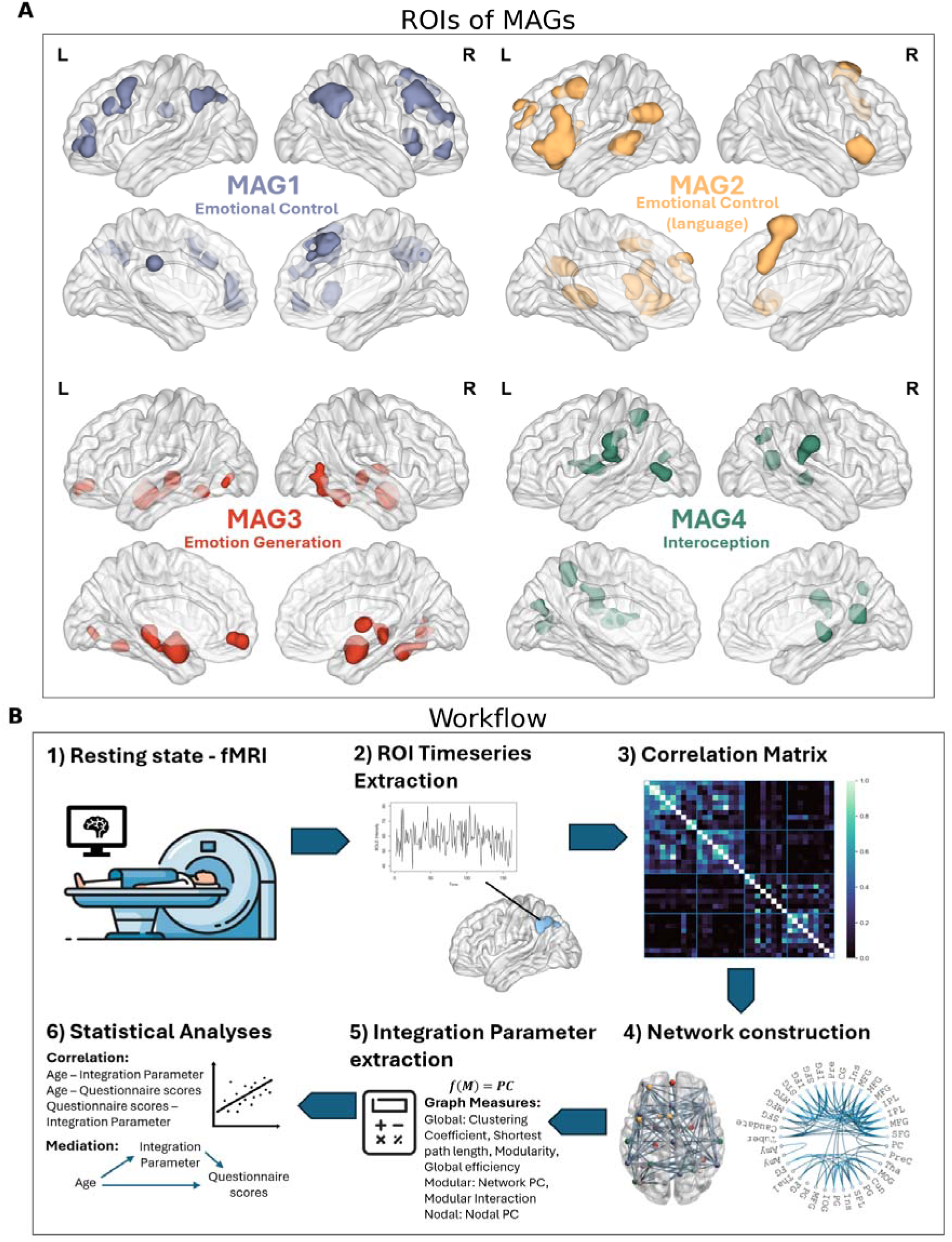
Visualization of Workflow with ROIs of MAGs: A) The ROIs of each MAG. MAG1 (indicated in purple) consisted of the bilateral dorsolateral prefrontal cortex (dlPFC) and inferior parietal lobule, right insula, cingulate gyrus, and precuneus. MAG2 (indicated in yellow) involved the bilateral ventrolateral prefrontal cortex (vmPFC), left temporoparietal junction, and middle temporal gyrus, and left dlPFC. MAG3 (indicated in red) was based on the bilateral amygdala and fusiform gyrus, left parahippocampal gyrus, periaqueductal grey, and vmPFC. MAG4 (indicated in green) consisted of the left superior parietal lobule, bilateral postcentral gyrus, left insula, precuneus, thalamus and posterior cingulate cortex. Exact coordinates and volumes are presented in Table 2. B) Flowchart of procedure 1) Participants were scanned at rest. 2) ROIs were masked, and their time courses extracted. 3) Time courses of all 36 ROIs of the MAGs were correlated with one another. 4) Graphs were derived from the correlation matrices with a sparsity of 0.3. 5) Mathematical measures were derived from the graph, including Network participation coefficient (PC), nodal PC, and network interactions, clustering coefficient, shortest path length, global efficiency and modularity. 6) Statistic analyses were performed on the graph measures, age, and behavior.

To test our first two hypotheses, that functional integration is increased with increasing age on the network and nodal level, participation coefficient (PC) and modular interaction were applied. The PC quantifies the relative connectivity of a node in its respective module compared to the entire graph.

With the PC of node i being *PC_i_ = 1 - Σ_m=1_^M^(k_im_/k_i_)*, m corresponds to the respective module (in our case M = 4, given the four MAGs), ki represents the total amount of edges a node has, and kim represents the count of edges the node has within its own module (Sporns et al., 2007). That is, if all the node’s edges are limited to its module, its participation coefficient is 0. Due to the sparsity of the graph, PC values will mostly be either 0 or above 0.5, because when the number of total edges is low, few connections outside the associated module will bring the PC into the higher ranges of 0.5 or more (Bloznelis, 2013).

To test the integration of the MAGs (Hypothesis 1) we calculated the modular PC by averaging the nodal PC values of each module. In addition, to address Hypothesis 1, we computed the modular interaction as the ratio of observed connections to the total possible connections. Due to the sparsity of the graph, the correlations between the time courses of certain ROIs did not exceed the threshold and were therefore not represented as an edge in the graph. Only edges that exceeded the sparsity threshold of 0.3 are considered “observed connections”. Thus, a modular interaction value of one represents the state that every possible connection that one module can make either within itself or with another module is also observed. To address which nodes of the network are more integrated in higher age and thus contribute to differences in network integration (Hypothesis 2), nodal PC was used.

To strengthen our analyses, we further examined clustering coefficient, shortest path length, global efficiency, and modularity, considering the four MAGs as one network, with the individual MAGs as modules (also reported in Figure S3). The clustering coefficient measures how many triangles a node can make via two of its neighbors, thus it approximates local segregation (Bloznelis, 2013; Tononi et al., 1998). The shortest path length reflects the average steps it needs from one node to another. Global efficiency is closely related to the shortest path length and serves as an estimation of how fast information can travel through the network (Seguin et al., 2023). However, in modular networks these measurements might not sufficiently reflect the structure of the network as the integration in the modules might differ from the integration of the whole network (Sporns & Betzel, 2016). To address the modular structure of the MAGs, we included a modularity measurement (Q), which decreasing modularity reflects how the network loses its modular structure and becomes more integrated overall. It also has to be noted that we distinguished whole-graph from module-level metrics. Modularity Q, clustering coefficient, shortest path length, global efficiency was computed once on the full 36-ROI graph. In contrast, participation coefficient (PC) and modular interaction were computed for each MAG and for each node; we did not average PC or interaction values across MAGs. Thus, all MAG-level results (e.g., “integration of MAG1”) refer to module-specific measures, not to graph-wide averages.

### 2.4 Network properties and association with age and behavior

To test our first hypothesis regarding age-related differences in the functional integration of the MAGs, partial correlations between age and modular PC for each module, as well as correlations between age and each of the modular interaction values were performed. For the second hypothesis regarding age-related differences in the functional integration of the ROIs in the MAGs, partial correlations between age and nodal PC for each node were performed. During these analyses, the effects of years of education, gender, and head motion were controlled for as partial regressors. Since in large samples, common normality tests like the Shapiro Wilk test often fail we assessed normality of each variable visually (Royston, 1982; Shapiro & Wilk, 1965). After visually assessing each variable’s normality, we determined whether parametric Pearson correlation or nonparametric Spearman correlation will be performed (to see variables distributions see **Figure S2**). For this we z-scored the data. We applied Bonferroni correction for multiple comparisons with α_adjusted_ = (0.05/ (4 modules + 36 nodes + 10 interactions)) = 0.001. To assess the relationship of age to network integration we fitted a generalized additive model to the data to address the possibility of nonlinearity in the data. Generalized Additive Modeling (GAM) is a method that incorporates smooth functions to represent the influence of variables. These functions can exhibit nonlinearity, depending on the inherent data patterns (Hastie, Trevor J., 1992). When the model involves nonlinear effects, such as possibly the impact of age on network integration, GAM enables to detect and incorporate typical nonlinear patterns that a traditional linear model might overlook. Our model contained one explanatory variable, age, which is captured by 7 penalized basis splines. Gender, years of education, and head movement were treated as linear regressor terms using Gaussian regression.

Additionally, for our third hypothesis regarding the association between network integration and psychological measures, we examined whether there was an age-related effect on any of the psychometric tests using partial correlation with subsequent linear regression. To assess whether the integration of specific nodes or modules had any associations with alterations in psychometric scores, we correlated all graph-derived variables with relevant psychometric score variables. To further explore whether age-related differences in behavior might be mediated by differences in MAG functional integration, mediation analyses were performed only for variable pairs that showed significant partial correlations in the initial analyses. Specifically, age was significantly associated with stress and loneliness, but not with depression or anxiety; therefore, mediation models were tested only for these significant associations. In total, three mediation models were examined, and the correction for multiple comparisons was applied across these three tests. Thus, for the significance value we chose α_adjusted_ = 0.05/3 = 0.017. In these mediation analyses, age served as the independent variable, the psychometric score as the dependent variable, and the network integration variable as the mediator. All variables were z-scored for this analysis. All graph theoretical analyses were carried out in the MATLAB-based toolbox GRETNA (Wang et al., 2015). All correlation analyses were performed using Python 3.9.16 with the packages SciPy.stats, statsmodels (Seabold & Perktold, 2010), and pingouin (Seongho Kim, 2015). Visualization was carried out using Python’s Seaborn and MATLAB’s BrainNetViewer (Xia et al., 2013). Mediation analysis was conducted using python statsmodels.

In order to assess if cognitive measurements impacted our findings, we simplified the complex cognitive measures from HCP-Aging data, Principal Component Analysis (PCA) of the scikit-learn package (sklearn.decomposition) in Python with two factors, accounting for 51.05% of the total variance, with the first and second component explaining 15.62%.

### 2.5 Validation

To further ensure the robustness and generalizability of the results we performed all the same analyses on resting-state fMRI data from the Cam-CAN for validation purposes, thereby confirming that the conclusions drawn are not due to peculiarities or biases in a single dataset but are indeed reflective of broader, reliable trends. The participant age ranges in Cam-CAN slightly differed from those in HCP-Aging, with some participants being younger than the youngest participant in HCP-Aging. To ensure consistency, the age range in Cam-CAN was adjusted to match that of HCP-Aging, by excluding those participants in the cam-CAN dataset that were younger than the youngest participant in the HCP-Aging dataset (HCP-Aging data set starts at 36 years old). Thus, analyses were performed on 333 subjects between the ages from 36 to 88. Additional preprocessing was carried out with the CONN default preprocessing pipeline which is identical to the processing of the HCP-Aging data. The parameters for graph analysis remained the same, and the covariates used in the regressions were also identical. Since Cam-CAN and HCP-Aging utilized different questionnaires, we used the behavioral scales in Cam-CAN that most closely resembled the psychometric data from HCP-Aging. In this instance, we found that the Hospital Anxiety and Depression Scales (HADSs) were the ones that captured the most alike aspects of mood in terms of the behavioral scales used in HCP-Aging (Harms et al., 2018; Hodes et al., 2013).

## 3 Results

### 3.1 Higher integration of MAG1 and lower integration of MAG4 in older age

#### 3.1.1 Increased global integration of MAG networks

We first characterized whole-graph properties (clustering, modularity) and then tested module-specific integration. We further assessed any differences in the integration of the MAGs across age, taking the four MAGs together as one large-scale network. Our results show the integration of all MAGs. Neither the shortest path length nor the global efficiency showed any significant differences between age groups. However, the clustering coefficient showed a positive association with age (*r* = 0.11, *p* = 0.009). Upon assessing the slope of the GAM fit, the increase of clustering seems to be linear with age. At the same time the modularity of the MAGs decreases with age (*r* = −0.22, *p* < 0.001). Taken together these findings indicate that the MAGs become more integrated with one another, losing their specificity (See Figure S3). In other words, the whole graph lower modularity with age, indicating more global integration. Critically, these global trends did not imply homogeneous changes across emotion networks: MAG-specific analyses revealed selective age effects—increased PC for MAG1 (emotional control) and decreased PC for MAG4 (interoception), with no age effects for MAG2 or MAG3. These findings, however, are not informative about whether this integration affects any certain modules (specific MAGs). Thus, to investigate such integration property within and between MAGs specifically, we used the PC and modular interaction for further analyses.

#### 3.1.2 Network PC and age association

Confirming our first hypothesis, the emotional control network (MAG1) showed a positive association between age and network PC (*r* = 0.260, *p* < 0.001; Figure 2A). The GAM also revealed a linear trend in all the splines.

**Figure 2.**
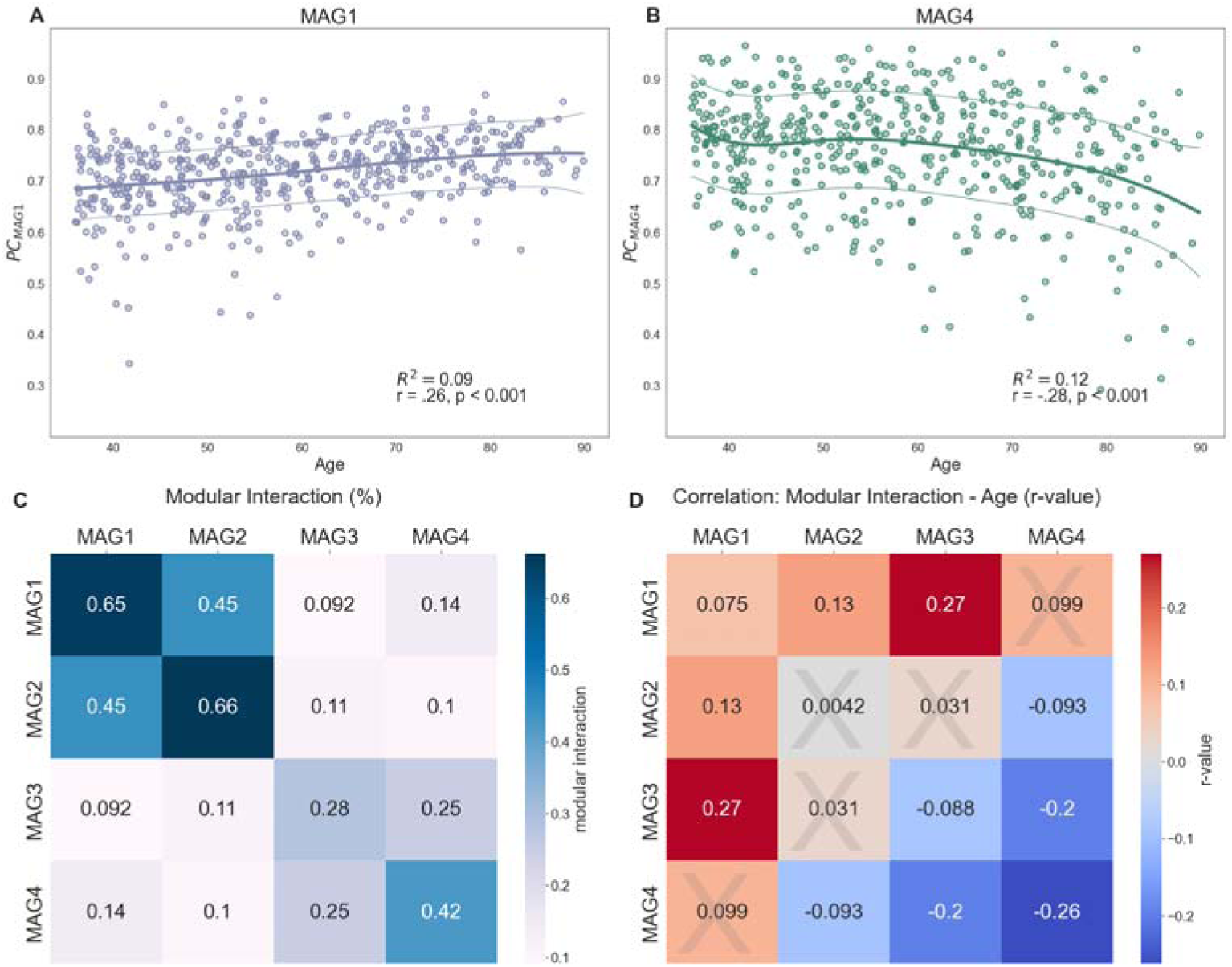
Functional integration of emotion regulation networks: (A) Generative additive model (GAM) fit of the integration of MAG1, the emotional control network, across age, showing a linear positive association between age and network PC. (B) GAM fit of network integration to age to MAG4, the interoception network. The model reveals a decrease of integration after the age of 70. (C) Average modular interaction between the MAGs in percent across the whole dataset, representing how many of all possible connections between nodes of the module are present. Darker color indicates high modular interaction. The modular interaction between the MAGs revealed a pattern of high connectivity between and within MAG1and MAG2, and MAG3 and MAG4, however showed low interaction between them. (D) Correlation values of this specific modular interaction with age. Blue fields indicate a negative association between modular interaction between these MAGs with age and red fields indicate a positive association between the interaction and age. Fields marked with a cross did not reveal a significant correlation with age.

On the other hand, we found age-related reduction in functional network integration of MAG4, which is a mediator between the network that is involved in both the perception of the emotional stimulus during the emotion generative process and during the regulation of emotional responses, that did not confirm our hypothesis. Rather, our data showed a significant shift towards lower PC of MAG4 over the time course of aging (*r* = −0.285, *p* < 0.001; Figure 2B). Together these findings partially support our first hypothesis, that brain networks involved in emotion regulation become more integrated in late adulthood. However, the GAM revealed that the integration of MAG4 followed a nonlinear pattern (pseudo *R*^2^ = 0.116), as the PC values were constant until the age of 70, followed by a steep decrease in network integration. This implies that only roughly after 70 MAG4 functional integration starts to decrease. No significant association was found between the PC of MAG2 and MAG3 with age (MAG2: *r* = −0.051, *p* = 0.207. MAG3: *r* = −0.042, *p* = 0.293).

#### 3.1.2 Age-related differences between MAGs are specific to MAGs 1 and 4

To further investigate Hypothesis 1, we examined the modular interaction between and within modules, which describes how many of the possible connections between all areas of one module to another are present. Modular interaction of one means all areas of both networks are connected, and zero means no functional connection. MAGs 1 and 2 seem to share most of the connections with one another in a functional community, and so do MAGs 3 and 4. However the interaction between the two communities (i.e., MAG1&2 with MAG3&4) is low (Figure 2C). Increasing integration of MAG1 with age was reflected in both increased within- and between-networks connections (Figure 2D). In the case of MAG1, this was indicated by the significant positive association between the modular interaction between MAG1 and both MAG2 (*r* = 0.208, *p* < 0.001) and MAG3 (*r* = 0.315, *p* < 0.001) and within itself (*r* = 0.154, *p* < 0.001) in later age. On the other hand, the decreased integration of MAG4 with age was mainly carried by a loss of connections to MAG2 (*r* = −0.175, *p* < 0.001) and MAG3 (*r* = −0.242, *p* < 0.001) and within itself (*r* = −0.302, *p* < 0.001). Further, MAG3 showed lower modular interaction within itself (*r* = −0.146, *p* < 0.001).

### 3.2 Specific nodes within a network contribute to overall network integration

To address Hypothesis 2, in which ROIs of the MAGs differ in functional integration during aging, we investigated the association of the integration of each individual ROI with age. The nodal PC of each node of the graph from each MAG was correlated with age.

We found that in MAG1, five ROIs showed a positive association between nodal PC and age (Figure 3). These ROIs included the bilateral middle frontal gyrus (right: *r* = 0,150, *p* < 0.001; left: *r* = 0.155, *p* < 0.001), left inferior parietal lobule (*r* = 0.165, *p*<0.001), right cingulate gyrus (*r* = 0.288, *p* < 0.001), and right precuneus (*r* = 0.157, *p* < 0.001). All these nodes showed a significant positive correlation between PC and age, indicating that these areas became more integrated globally with the other MAGs with increasing age.

**Figure 3.**
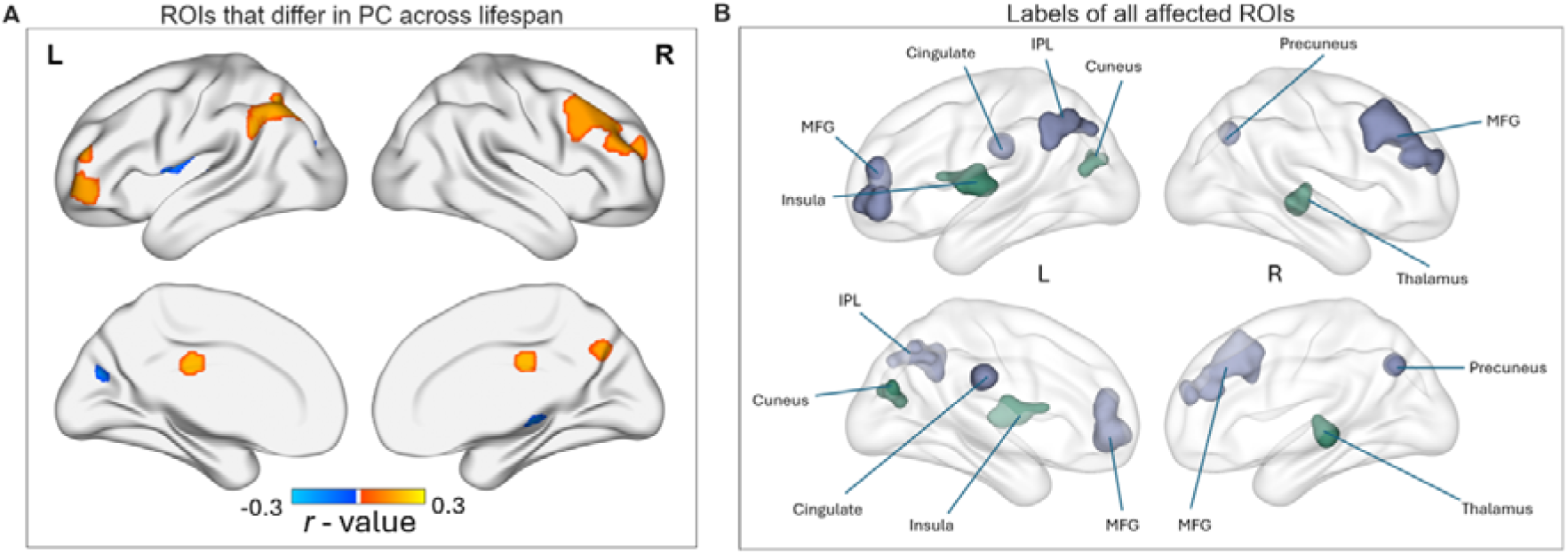
Functional integration of specific brain areas associated with MAGs in aging: (A) the r values for the ROIs that showed significant Pearson correlation with age demonstrated on the brain. Orange hues represent the areas that are more integrated in older adults while blue color indicate the areas that are less integrated with age. The areas that show increased integration correspond to the bilateral middle frontal gyrus, left inferior parietal lobule, right cingulate gyrus, and right precuneus, while the areas that show decreased integration correspond to the left insula, left cuneus and right thalamus. (B) Glass-brain showing also the medial areas which show altered integration across the aging lifespan and their network association (MAG1: purple, MAG4: green), p<0.001, Bonferroni corrected.

A negative association between PC and age in three ROIs in MAG4 (Figure 3) were found in the left insula (*r* = −0.198, *p* < 0.001), left cuneus (*r* = −0.157, *p* < 0.001), as well as right thalamus (*r* = −0.169, *p* < 0.001; see also **Figure S4** for further results from different brain nodes). These areas were less integrated in higher age, suggesting that they made proportionally less connections with the other MAGs with increasing age. No other significant correlations were found between age and any of the nodes from other MAGs. These findings altogether indicate that frontal ROIs showed positive association between nodal PC and age, more integrated with other MAGs with age, while there are also medial and lateral areas affected, becoming less integrated with age, confirming our second hypothesis.

### 3.3 Inter-MAG1 connectivity is associated with increased depression while integration of the left cuneus suppresses age effect on loneliness and perceived stress

To test the third hypothesis that differences in emotional regulation could be attributed to differences in emotion network integration, partial correlations with the integration parameters of the nodes and modules and psychometric scores were performed. We found that the within modular interaction of MAG1 correlated positively with depression scores (*r* = 0.142, *p* < 0.001), indicating that across different ages, more functional connections between the ROIs of MAG1 indicate a higher tendency towards depressive behaviors (**Figure S5**).

To further test whether alterations in network integration could explain age-related mental-wellbeing changes, mediation analyses was conducted. First, linear associations between psychological questionnaires and age were assessed, to examine the association between the independent variable (i.e., age) and the dependent one (i.e., psychometric scores). Self-report questionnaires provided by the HCP-Aging included measures of anxiety, depression, stress, and loneliness. Of these, perceived stress and loneliness (Figure 4A) showed significant decrease with increasing age. Then we performed linear regression between functional integration of the MAGs as measured by modular/nodal PC and modular interaction, with psychometric scores, to check which network topographical index could constitute a mediating variable.

**Figure 4.**
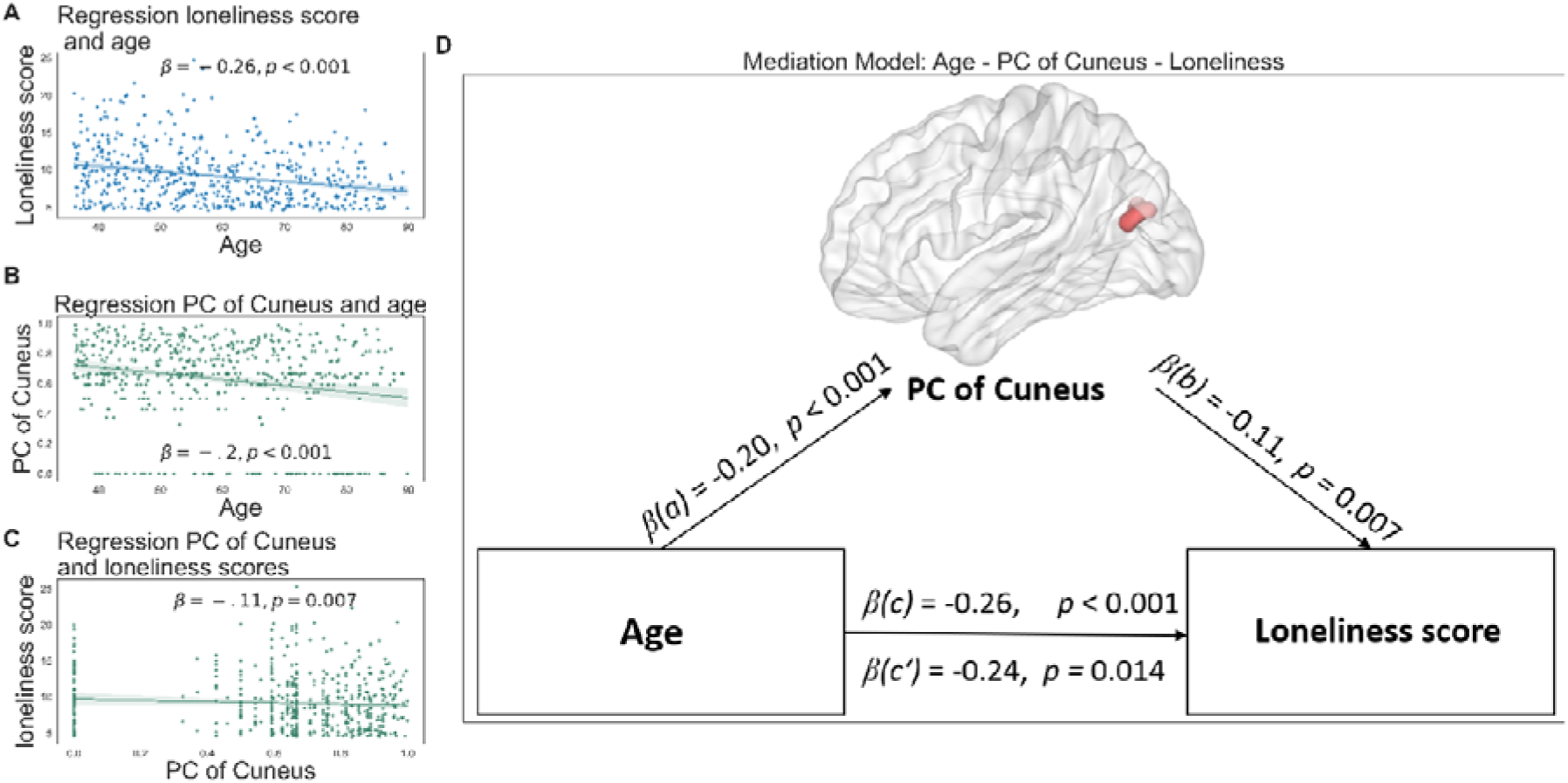
Association between age, cuneus functional integration and perceived stress and loneliness in the HCP-Aging dataset. (A) Linear regression between age and loneliness scores showing negative association with age. (B) Linear regression between functional integration of the left cuneus and age, showing lower PC with higher age. (C) Linear regression between functional integration of the left cuneus and loneliness scores, indicating a small negative trend. (D) Mediation model between age and loneliness scores, with cuneus PC mediating the effect partially suppressing the negative age effect on perceived stress. Β(c) = direct effect and β(c’) = indirect effect. The area marked in red are the respective voxels of the cuneus.

For the mediation analyses, age was used as the independent variable, perceived stress and loneliness as dependent variables in two separate mediations and network integration measures from all MAGS as mediators. Only one mediation model was significant, in the case of which it was a partial suppression (Smith et al., 1992). A suppressor variable does not have a direct effect on the dependent variable but, when included in the analysis, increases the relationship between the independent variable and the dependent variable. (for review on suppressor variables in psychology, see (Rucker et al., 2011). The direct effect from age on loneliness was negative (β = −263, *p* < 0.001) with the total effect (via cuneus integration) was slightly less negative (β = −0.240, *p* = 0.014), suggesting that the integration of the left cuneus of MAG4 suppressed the negative age effect on loneliness scores. Hence, for those older individuals that showed rather high cuneus integration even in older age, also exhibited increased loneliness scores. Thus, decreased PC of the cuneus suppressed the negative association of age and loneliness (Figure 4D). Since there was no other significant correlation between the integration of the MAGs and other psychometric scores, those mediation models were not considered (for all mediation models see **Table S2**). It should be noted that the results of mediation analysis with loneliness scores did not change following including cognitive PC scores into the model.

### 3.4 Validation with Cam-CAN dataset confirms increased integration of MAG1

Following the same analyses steps as for the analysis on the HCP-Aging dataset, we found that the findings on the HCP-Aging could be partially validated by the Cam-CAN data. Regarding our first hypothesis, a positive association of the PC of MAG1 with age, in the Cam-CAN dataset was confirmed (*r* = 0.10, *p* = 0.029, **Figure S6**). Other findings regarding the nodal changes from each MAG were confirmed in the Cam-CAN data including increased PC of the Inferior Parietal Lobule of MAG1 (*r* = 0.13, *p* = 0.004) in later age, decreased modular interaction of connections between MAGs 3 and 4 (*r* = −0.096, *p* = 0.037).

As for the second hypothesis, a measure of positive or negative affect in later age Cam-CAN provided the HADS depression and anxiety scores of the subjects. We found a negative correlation between HADS anxiety scores and age (*r* = −0.185, *p* < 0.001). This might reflect the behavioral findings of decreased stress and loneliness in the HCP-Aging.

None of the findings in HCP-Aging were contradicted by the Cam-CAN data, however some associations between integration measures of the MAGs and age present in the HCP-Aging data were not significant in the Cam-CAN dataset and vice versa. There were additional significant associations between network integration and age. Details of these differences are presented in the supplementary table (**Table S3**). All nonlinear models of Cam-CAN and HCP-Aging in all MAGs are presented in **Figure S6**.

## 4 Discussion

In the present study, our objective was to explore age-related alterations in the functional network architecture of emotion regulation networks applying graph network analyses to four major networks involved in emotion regulation and generation systems. Identifying how network topology might alter across the lifespan provides valuable insights into how a shift in the emotional wellbeing can occur in aging. We have gathered evidence indicating that across the adult lifespan alterations in large-scale brain networks associated with the emotional response occur, which correlate to differences in psychological well-being in later age that are discussed in detail below.

Although aging is associated with broad reductions in network segregation, our design isolates emotion-related modules (MAGs) derived from meta-analytic task evidence. Within this constrained functional space, age effects were not global: only MAG1 showed increased integration with age and only MAG4 showed reduced integration (with a late-life inflection), while MAG2/MAG3 were unchanged. Node-level effects likewise clustered within frontal control nodes (MAG1) and interoceptive/sensory nodes (MAG4). Behavior–brain associations were also network-specific (e.g., within-MAG1 interaction and depression; MAG4 cuneus integration and loneliness). Replication of MAG1 findings in an independent dataset (Cam-CAN) further supports domain specificity within emotion-related systems. Thus, the observed age effects are not merely general aging signatures but selective alterations in emotion regulation networks.

### Increased integration of emotion control network

Partially confirming our hypothesis, we observed an increase in functional integration of the emotional control network, namely MAG1, with increasing age. We also observe decreased integration of interoception network, namely MAG 4, in late adulthood. Interestingly the regions in MAG1 that show significant positive association with age are areas involved in high-order function, while the significant regions in MAG4 are except the insula, mostly sensory areas (Huntenburg et al., 2018). Network integration has previously been debated in the context of balancing metabolic costs and communication efficiency (Hillary & Grafman, 2017). Thus, specifically in the older brain that is faced with metabolic changes (Mattson & Arumugam, 2018), increased integration might reflect a compensatory mechanism for metabolic changes. More generally, a recent study has shown that better crystallized intelligence is associated with higher global integration, while on the other hand, processing speed and memory recruitment are associated with lower brain integration. While the mechanisms of why this network alteration occurs is still unknown, the current study was able to show alteration in the brain organization in the emotion regulation networks across the lifespan. Perhaps the observed increased integration of MAG1 might reflect regulatory control in later age over peripheral and arousal systems (Mather, 2024), which require higher metabolic demand. Similarly, decreased integration of MAG4 could reflect a conservation of processing speed during emotion generation and reflection. A fast emotional response is crucial to individuals irrespective of age, therefore this function might be preserved into higher ages, reflected by the decreased integration of MAG4. Although the picture of emotion regulation in aging is quite nuanced. Our interpretation of the increased integration of MAG 1 with age does not imply that older adults are more effective at emotion regulation per se. Indeed, older adults are not necessarily better at employing cognitively demanding strategies such as reappraisal (Livingstone & Isaacowitz, 2019). Rather, we suggest that this pattern reflects altered recruitment of prefrontal regions implicated in regulatory and evaluative processes, which may support the maintenance of emotional well-being despite age-related neural changes. In other words, increased MAG 1 integration may serve to preserve adaptive emotional outcomes rather than to enhance regulatory efficiency itself. This interpretation aligns with prior findings showing that interoceptive awareness declines with age (Khalsa et al., 2009), suggesting that reduced communication between interoceptive and higher-order cortical regions may underlie diminished internal bodily awareness in older adults.

The MAG1 consists of mostly frontal regions, including the superior and middle frontal gyrus. This finding aligns with previous studies that suggest the increased prefrontal recruitment in emotion regulation tasks that is observed in older adults (Berboth & Morawetz, 2021; Nashiro et al., 2012). In line with recent meta-analytic evidence, MacCormack et al. (2020) reported that older adults show greater functional co-activation in frontal control regions, whereas younger adults exhibit stronger co-activation in amygdala and midcingulate cortices during affective tasks. These findings converge with the present results, suggesting a broader shift with age from limbic to prefrontal involvement in emotional processing. Particularly the bilateral MFG, left IPL, precuneus, and cingulate nodes of this network showed increased integration in later life, all of which have been demonstrated to be crucial in the regulation of emotions and for emotional wellbeing and stability (Glinka et al., 2020; Tomasino et al., 2022; Williams et al., 2006; Winecoff et al., 2011; Ziaei et al., 2017, 2021). On top of that, MAG1 is predominantly involved in the downregulation of negative emotions (Morawetz et al., 2020). We suggest that as MAG1 becomes more integrated in later life, there is an enhanced transmission of information to the other networks (Sporns, 2013) that are involved in generating and processing emotional responses, which in turn could have beneficial effects on mood and emotional well-being in late life. Although not directly tested in this study, our findings support the socioemotional selectivity theory and may suggest that increased functional integration of the MAG1, or emotional control network, can contribute to the positivity effect observed in late adulthood. Many of the ROIs belonging to MAG1 are also associated with other large-scale brain networks such as the DMN or frontoparietal network (FPN). It is specifically these areas of MAG1 that also exhibited increased nodal PC in higher age, including the MFG, inferior parietal lobule and cingulate gyrus. Thus, the observed increased integration of MAG1 in the current study is consistent with previous findings that there is increased network integration of the DMN and FPN in aging (Andrews-Hanna et al., 2007; Ng et al., 2016). In a study investigating the test-retest reliability of the MAGs at ultrahigh magnetic field strength, the first two MAGs showed high to excellent reliability in the context of emotion regulation tasks (Berboth et al., 2021), further supporting our findings.

Moreover, we found that there is a positive association between the inter-module interaction of MAG1 and depression scores, regardless of the individual’s age. Indeed, a depressive phenotype has previously been associated with an increased top-down processing from the prefrontal areas towards limbic areas (Challis & Berton, 2015; LeDuke et al., 2023; Mogg & Bradley, 2016). MAG1 consists mostly of frontal areas and exhibits a rather top-down emotional control function (Morawetz et al., 2020). We showed that the number of connections that MAG1 makes within itself is higher in subjects with higher depression scores. This finding could reflect a tendency towards increased top-down inhibition as many prefrontal areas that are part of MAG1 are more functionally connected. Moreover, since MAG1 shares several regions with the DMN, this finding might reflect the observed increased functional connectivity of the DMN in major depression (Gillespie et al., 2020; Bao-Juan Li et al., 2018). Our current finding replicates the involvement of the functional connectivity increase of certain nodes within the DMN in depression and extends it with the other MAG1 areas that are not part of the DMN to contribute to the altered functional connectivity that is involved in the neuropathy of depression.

It remains critically important to investigate the underlying drivers of changes in functional network integration, as well as to apply other methodological advances to further examine age-related changes or differences in topological properties. Potential candidates for investigation include changes in receptor expression for different transmitters (Kringelbach et al., 2020; Shine, 2019), or differences in white matter development (Stevens et al., 2009). Our findings underscore the significance of functional network integration in emotional wellbeing across the adult lifespan. However, with a deeper understanding of what precipitates these changes, it could become a target for therapeutic intervention aimed at enhancing emotional wellbeing in later life. Although the current study is correlational, the observed age-related changes in functional integration may help identify neural systems that could serve as targets for future therapeutic approaches. For example, interventions that enhance prefrontal–control network function (e.g., cognitive–emotional training, mindfulness-based programs) may strengthen the adaptive integration patterns observed in MAG1, whereas approaches that foster awareness of bodily states (e.g., interoceptive training, biofeedback, or gentle physical practices) could help maintain communication between interoceptive and cortical regions such as the cuneus and insula (MAG4). In this context, therapeutic interventions aimed at enhancing emotional wellbeing refers to strategies designed to promote emotional balance and resilience in aging, rather than specific treatments directly derived from the present findings. In addition, recent hierarchical gradient analyses of the frontoparietal network, also part of MAG1, confirmed our findings that it becomes more integrated or dedifferentiated with age, thereby validating our results through an independent advanced methodological approach (Ye et al., 2025).

### Decreased integration of interoception network in aging

We observed decreased integration of MAG4 after the age of 70, which has been identified to be involved both in emotion generation and regulatory processes, particularly linking the two networks via interoception. Interoception is vital to perceive our own body states and helps shape the feeling and intensity of emotions (Craig, 2002). Existing research, however, provides limited information on how age differences in interoception may affect emotional experience and general well-being. Our study shed some light on the fact that the decreased integration of MAG4 in older age may contribute to the decline in interoceptive capabilities. According to previous studies, some areas such as the insula, which is a critical area for interoception, decrease in volume with age (Good et al., 2001). We report significantly decreased integration of the left insula in late adulthood, which could be attributed to decay in volume and could further explain difficulties in interoceptive processes as part of emotion regulation networks in older adults. These results should be confirmed with targeted studies on interoception in the future studies combining volumetric measures with large-scale network architecture. It must be noted that the observed decrease in integration does not follow a fully linear trend, but the rate of change seems to increase only after the age of 70. Previous studies find accelerated white matter decay, after the age of 70 (Fjell & Walhovd, 2010; Resnick, 2000) which could contribute to decreased integration of MAG4. This age is also characterized as the start of a decline in interoceptive awareness, including awareness of visceral sensations such as esophageal pain, gastric distension, and heartbeats (Rayner, MacIntosh, 2000; Khalsa et al., 2009; Lasch et al., 1997). The brain areas within MAG4 that contributed the most to the overall decreased integration of this network included the insula, the cuneus, and the right thalamus. These areas have been linked with empathy and relating to another person’s negative emotion or pain (Bufalari et al., 2007; Nummenmaa et al., 2008; Ziaei et al., 2021). We speculate that lower integration of these areas and consequent reduced signaling from them may indicate older adults’ difficulty reading and processing negative affect in others (Hayes et al., 2020; Mill et al., 2009; Stutesman & Frye, 2023). However, it could also reflect a preservation of emotion and empathetic processes in later life. Taken together, the increased integration of nodes in MAG1 that are mostly associated with emotional control or cognitive control tasks (Friedman & Robbins, 2022; Koch et al., 2018) with the decreased integration of nodes in MAG4 that are associated with empathy and relating to others (Arioli et al., 2021; Braadbaart et al., 2014) could reflect that emotion processing in older adults relies more on crystalline aspects of cognition like knowledge about oneself and the world. A process that has previously been described in the cognitive domain as the crystallization of cognition (for review see Spreng and Turner, 2019) and has also been suggested to affect the sociocognitive domain (Henry et al., 2023). The dual pattern of higher integration of the frontal emotion regulation network (i.e. MAG1) with lower integration of the mediating emotion regulation and generation network (i.e. MAG4) in later age could reflect that not all aspects of emotional processes and social cognition are affected at the same level and rate in late adulthood. While control over negative emotions is preserved or even enhanced with age, other aspects, such as interoception, may be more adversely affected. This finding aligns with prior research and reflects the dual association observed through topological measures in this study. Future research should explore whether age-related structural changes in specific nodes of MAG1 and MAG4, or differential receptor expression within these networks, contribute to these dual association patterns.

Taken together, it is important to note that network integration and within-network modular interaction represent distinct aspects of functional organization. Integration captures the extent to which a given module communicates with other large-scale systems across the brain, whereas within-network modular interaction reflects internal coherence among nodes belonging to the same module. Thus, a network may be highly integrated with the rest of the brain while exhibiting relatively low within-network coherence, and vice versa. Our hypotheses were formulated around integration, not within-network connectivity, as we aimed to examine how large-scale communication between emotion-related modules and the broader functional architecture of the brain relates to behavioral outcomes in aging. The observed relationship between greater within-network coherence of MAG1 and depressive symptoms was therefore not anticipated, but may suggest that increased internal coupling within prefrontal–control regions could reflect maladaptive or rigid emotional control processes, such as rumination, which are commonly associated with depressive states.

In contrast, the reduced integration of MAG4 observed in older adults likely reflects decreased communication between interoceptive regions and other large-scale systems. While this can indeed be viewed as reduced inter-network communication, our interpretation considers that such reduced integration may also represent a compensatory mechanism—preserving efficient, streamlined processing for emotionally salient information in the context of aging. These interpretations should be considered tentative and future research is needed to further examine the causal link between these constructs.

The question remains whether the observed changes in network integration relate to improved mood in late adulthood. In the HCP-Aging dataset we found a decline in perceived stress as well as loneliness in later life, therefore, we were interested if one of the observed differences in brain network integration could mediate this effect. There was only one significant mediation model suggesting that the decreased integration of the left cuneus as a partial suppressor (Yongnam Kim, 2019) reverts age-related decrease in perceived loneliness. While the cuneus is mostly regarded as a visual area, it has also been associated with sensory integration and psychiatric conditions (Martel et al., 2017; Nyatega et al., 2021). Activity and volume of the left cuneus has been associated with symptom severity in depression and anxiety (Brühl et al., 2014; Dotson et al., 2022; Guo et al., 2012; Peng et al., 2019; Yoon et al., 2016) and the ability to adjust one’s emotion (Tan et al., 2014) In healthy aging the activity of the visual and parietal areas tends to decrease (Springer et al., 2023). One could speculate that the Cuneus constitutes an important relay area integrating visual information (Xiaoxi He et al., 2021; Nyatega et al., 2021). Thus, this finding might indicate that the age-related decrease of cuneus integration might alter brain networks’ functional connectivity in a way that is beneficial on the perception of one’s own loneliness in later life. In other words, these findings suggest that the observed association between cuneus integration and loneliness may reflect differences in internally generated representations of social and emotional experiences. In this context, reduced integration of the cuneus could indicate less efficient communication between perceptual and self-referential systems, potentially contributing to altered emotional appraisal or diminished social connectedness in aging

### Integration of emotion regulation network across datasets

To ensure the generalizability and reproducibility of our results, a critical step in our methodology involved the validation of our key findings from the HCP-Aging using the Cam-CAN dataset. Our main finding, the increased integration of the frontal emotional control network, was replicated in the Cam-CAN dataset. However, the decrease in integration of the interoceptive network, MAG4, did not survive in the Cam-CAN dataset. One reason for such difference between the two datasets could be attributable to different scanning protocols (including different scanner systems and different repetition times, **Table S1**), difference in the duration of scan times and having participants closing their eyes or not (Han et al., 2023; L. Wu et al., 2010). The replication of the finding that MAG1 is more integrated in later life in eyes-open and eyes-closed resting state supports that this finding is a general trend which is not condition-dependent. Another aspect that must be considered in the nature of the two datasets is that the countries of origin of the data are different, which is important while studying social-emotional processes across the lifespan that might be affecting these findings (Foulkes & Blakemore, 2018; Kivimäki et al., 2020; Tooley et al., 2021).

### 4.1 Limitations and future perspectives

Due to the nature of the method using mathematical graphs, the interpretation of the findings needs to be carefully evaluated. An increase in the integration of a functional network might indicate increased information transmission from one network to another but could also imply functional blurring and the loss of specialized processing in the network (Sporns, 2013). In the context of aging, the increased global integration across the whole brain has been associated with decreased performance in cognitive demands tasks (Pedersen et al., 2021). Hence it cannot be concluded whether the increase in MAG1 integration in later age is in any way beneficial or detrimental for the whole system. A conservation of the low integration of other MAGs in later life might therefore also be an indicator as to why emotional processes are preserved longer in later life even though other cognitive processes decline. Another limitation that would also count as an advantage is the size of the HCP-Aging and Cam-CAN datasets. The large sizes of the data allow also small effects to be found, however, our findings remain to be replicated in future studies using small samples. Our findings yield a great starting point for further research to explore the role of emotion regulation networks not only in aging but across all ages. Future studies should also consider applying model-based methods such as independent component analysis to investigate shared or unique networks with the model-based method using MAGs here. Furthermore, future research could investigate whether these networks change across the lifespan occur in longitudinal studies, employing similar or additional graph measurements to investigate the functional topology of these networks across lifespan and their changes with age.

### 4.2 Conclusion

In the present study, we found alterations in the functional architecture of emotion regulation networks across the adult lifespan. The emotional control network (i.e., MAG1) became more integrated with the other networks, consistent across two large datasets, the HCP-Aging, and Cam-CAN. Additionally, we discovered that the network mediating emotion regulation and generation, the interoception network (i.e., MAG4), became more segregated. To link these findings to emotional wellbeing in late adulthood we identified a mediating role on loneliness of the cuneus, an area that only recently is understood to be more than a visual area but has impacts on more abstract cognitive functions. Overall, identifying topological changes in emotion network architecture is of particular interest as it allows for further understanding of emotional aging and late-life mood disorders.

## Disclosure

The authors declare no competing interest.

## Availability of data and code

Code used for this project is available through Open Source Framework: <u>OSF | Aging, Emotion</u> <u>regulation, Brain networks</u>

## Funding

This project was funded by Kavli Foundation and was partially supported by the Research Council of Norway through its Centers of Excellence scheme, project number 332640.

## CRediT authorship contribution statement

**Leona R. Bätz:** Writing – original draft, Writing – review & editing, Methodology, Investigation, Formal analysis, Data curation, Visualization, Validation. **Shuer Ye:** Writing – review & editing, Software, Supervision, Methodology, Data curation, Resources, Methodology, Investigation, Conceptualization. **Xiaqing Lan:** Writing – review & editing, Resources, Data curation. Melissa Schubbert: Writing – review & editing, Analysis**. Maryam Ziaei:** Writing – review & editing, Supervision, Project administration, Conceptualization, Funding acquisition.

## Abbreviations

PFC: Prefrontal Cortex
MAGs: meta-analytic groupings
fMRI: functional magnetic resonance imaging
HCP: Human Connectome Project
Cam-CAN: Cambridge Center for Aging and Neuroscience
PC: participation coefficient
GAM: General additive model
DMN: default mode network

## Acknowledgements

Research reported in this publication was supported by the National Institute on Aging of the National Institutes of Health under Award Number U01AG052564 and by funds provided by the McDonnell Center for Systems Neuroscience at Washington University in St. Louis. The HCP-Aging 2.0 Release data used in this report came from DOI: 10.15154/1520707.

Data collection and sharing for this project was provided by the Cambridge Centre for Ageing and Neuroscience (CamCAN). CamCAN funding was provided by the UK Biotechnology and Biological Sciences Research Council (grant number BB/H008217/1), together with support from the UK Medical Research Council and University of Cambridge, UK. This work was partially supported by the Research Council of Norway through its Centres of Excellence scheme, project number 332640.

## Notes

https://osf.io/npmeb/

### Competing Interest Statement

The authors have declared no competing interest.

### Summary of Updates

Revised following reviewers comments and now accepted in Aging Brain journal

