## supplementary for "Increased functional integration of emotional control network across the adult lifespan"

Abbreviated title: Emotion regulation networks across the lifespan

Authors: Leona Rahel Bätz^1*^, Shuer Ye^1^, Xiaqing Lan^1^, Melissa Schauber^1^, Maryam Ziaei^1,2*^

^1^ Kavli Institute for Systems Neuroscience, Norwegian University of Science and Technology, Trondheim, Norway

^2^ K.G. Jebsen Centre for Alzheimer's disease, Norwegian University of Science and Technology, Trondheim, Norway

*Corresponding Author2:

### **Supplementary Tables**

**Table S1: Comparison of BOLD and T1 scanning parameters employed to produce HCP-Aging and Cam-CAN datasets.**

| **BOLD Parameter** | **HCP-Aging** | **Cam-CAN** |
| --- | --- | --- |
| Scanner System | Siemens MAGNETOM Prisma | Siemens TIM TRIO |
| Field strength | 3 Tesla | 3 Tesla |
| Repetition time (TR) | 800 ms | 1970 ms |
| Time to Echo (TE) | 37 ms | 30 ms |
| Pulse Sequence | 2D multiband gradient EPI | T2* GE EPI |
| Voxel Size | 2×2×2 mm | 3×3×4.44 mm |
| Slices | 72 axial slices | 32 axial slices |
| Flip Angle | 52° | 78° |
| Paradigm | Eyes-open resting state | Eyes-close resting state |
| Scan duration | 6 minutes 5 seconds × 4sessions | 8 minutes 40 seconds |
| **T1 Parameter** | **HCP-Aging** | **Cam-CAN** |
| Scanner System | Siemens MAGNETOM Prisma | Siemens TIM TRIO |
| Field strength | 3 Tesla | 3 Tesla |
| Repetition time (TR) | 2500 ms | 2250 ms |
| Time to Echo (TE) | 1.8/3.6/5.4/7.2 ms | 2.99 ms |
| Inversion Time (TI) | 1000ms | 900 ms |
| Pulse Sequence | multi-echo MPRAGE | MPRAGE |
| Voxel Size | 0.8×0.8×0.8 mm | 1×1×1 mm |
| FOV | 256 × 240 × 166 mm | 256 × 240 × 192 mm |
| Flip Angle | 8° | 9° |
| Scan duration | 8 minutes 22 seconds | 4 minutes and 32 seconds |

**Table S2: beta values and significance for all mediation models.** All mediation models were Bonferroni corrected with α_corrected_=0.05/3 = 0.017, thus only the mediation between age, cuneus PC and loneliness survived the correction.

| **Model** | **β(X→Y)** | **p** | **β(X→M)** | **p** | **β(M→Y)** | **p** | **β(X→M→Y)** | **p** |
| --- | --- | --- | --- | --- | --- | --- | --- | --- |
| X: age,  Y: Stress,  M: Modular Interaction MAG1 | -0.2691 | <0.001 | 0.1550 | <0.001 | -0.0856 | 0.039 | -0.013345 | 0.072 |
| X: age,  Y: Stress,  M: Cuneus PC | 0.3007 | <0.001 | -0.1990 | <0.001 | -0.0920 | 0.027 | 0.018503 | 0.04 |
| X: age,  Y: Loneliness,  M: Cuneus PC | -0.2631 | <0.001 | -0.1990 | <0.001 | -0.1139 | 0.007 | 0.022944 | 0.014 |

**Table S3: Correlations between Network measure and age in the Cam-CAN and HCP-Aging datasets.** R values from Pearson correlation with their respective significance value, which were Bonferroni corrected for findings in the HCP dataset (p_corrected_ < (0.05/50) = 0.001; N= 4 Networks + 36 nodes + 10 modular interactions = 50).

|  | **HCP-Aging** | | **Cam-CAN** | |
| --- | --- | --- | --- | --- |
|  | *r* | *p* | *r* | *p* |
| ***Network PC*** |  |  |  |  |
| MAG1 | 0.23 | < 0.001 | 0.10 | 0.029 |
| MAG2 | -0.6 | 0.09 | -0.005 | 0.904 |
| MAG3 | -0.05 | 0.2 | 0.01 | 0.739 |
| MAG4 | -0.28 | < 0.001 | 0.25 | 0.05 |
| ***Nodal PC*** |  |  |  |  |
| Superior Frontal Gyrus | 0.0029 | 0.93 | 0.07 | 0.111 |
| Middel Frontal Gyrus | 0.138 | < 0.001 | 0.08 | 0.074 |
| Inferior parietal lobule | 0.143 | < 0.001 | 0.05 | 0.210 |
| Inferior Parietal Lobule | 0.168 | < 0.001 | 0.13 | 0.004 |
| Middle Frontal Gyrus | 0.103 | 0.006 | 0.02 | 0.671 |
| Middle Frontal Gyrus | 0.071 | 0.063 | 0.05 | 0.293 |
| Middle Frontal Gyrus | 0.1 | 0.008 | 0.15 | 0.001 |
| Insula | -0.026 | 0.497 | 0.002 | 0.956 |
| Cingulate Gyrus | 0.186 | < 0.001 | -0.07 | 0.087 |
| Precuneus | 0.151 | < 0.001 | 0.06 | 0.219 |
| Inferior Frontal Gyrus | 0.013 | 0.719 | 0.11 | 0.012 |
| Superior Frontal Gyrus | -0.042 | 0.263 | -0.10 | 0.024 |
| Inferior Frontal Gyrus | -0.06 | 0.112 | 0.03 | 0.392 |
| Superior Temporal Gyrus | 0.053 | 0.164 | -0.02 | 0.559 |
| Middle Temporal Gyrus | -0.063 | 0.095 | -0.01 | 0.863 |
| Middle Frontal Gyrus | -0.066 | 0.083 | 0.07 | 0.972 |
| Superior Frontal Gyrus | 0.032 | 0.392 | -0.036 | 0.559 |
| Caudate | -0.129 | 0.0006 | 0.064 | 0.891 |
| Tuber | 0.115 | 0.002 | -0.03 | 0.146 |
| Amygdala | -0.036 | 0.335 | -0.15 | 0.001 |
| Amygdala | -0.011 | 0.773 | 0.004 | 0.390 |
| Fusiform Gyrus | -0.062 | 0.101 | 0.08 | 0.068 |
| Thalamus | 0.053 | 0.16 | 0.098 | 0.070 |
| Fusiform Gyrus | -0.054 | 0.156 | 0.004 | 0.948 |
| Parahippocampal Gyrus | -0.04 | 0.289 | 0.07 | 0.193 |
| Medial Frontal Gyrus | -0.027 | 0.481 | 0.076 | 0.481 |
| Inferior Occipital Gyrus | -0.011 | 0.774 | -0.03 | 0.884 |
| Postcentral Gyrus | -0.188 | < 0.001 | 0.006 | 0.942 |
| Insula | -0.183 | < 0.001 | -0.03 | 0.100 |
| Superior Parietal Lobule | -0.146 | 0.0001 | 0.11 | 0.019 |
| Postcentral Gyrus | -0.171 | < 0.001 | 0.019 | 0.231 |
| Cuneus | -0.188 | < 0.001 | 0.018 | 0.067 |
| Middle Occipital Gyrus | -0.081 | 0.034 | 0.021 | 0.168 |
| Thalamus | -0.171 | < 0.001 | 0.002 | 0.959 |
| Precuneus | -0.039 | 0.302 | 0.03 | 0.457 |
| Posterior Cingulate | 0.007 | 0.849 | 0.03 | 0.415 |
| ***Modular Interactions*** |  |  |  |  |
| Within MAG1 | 0.165 | < 0.001 | -0.011 | 0.812 |
| Within MAG 2 | 0.06 | 0.112 | -0.120 | 0.009 |
| Within MAG 3 | -0.139 | < 0.001 | 0.044 | 0.339 |
| Within MAG 4 | -0.3 | < 0.001 | -0.034 | 0.459 |
| Between MAG1&2 | 0.212 | < 0.001 | 0.029 | 0.523 |
| Between MAG1&3 | 0.267 | < 0.001 | 0.071 | 0.122 |
| Between MAG1&4 | 0.026 | 0.483 | 0.134 | 0.003 |
| Between MAG2&3 | 0.022 | 0.555 | -0.040 | 0.387 |
| Between MAG2&4 | -0.189 | < 0.001 | -0.043 | 0.347 |
| Between MAG3&4 | -0.225 | < 0.001 | -0.096 | 0.037 |

Note. Green highlights indicate the main findings in HCP-Aging, yellow indicates the finding is an additional finding in Cam-CAM, and blue indicates main findings in HCP-Aging were validated in Cam-CAN.

**Supplementary Figures**

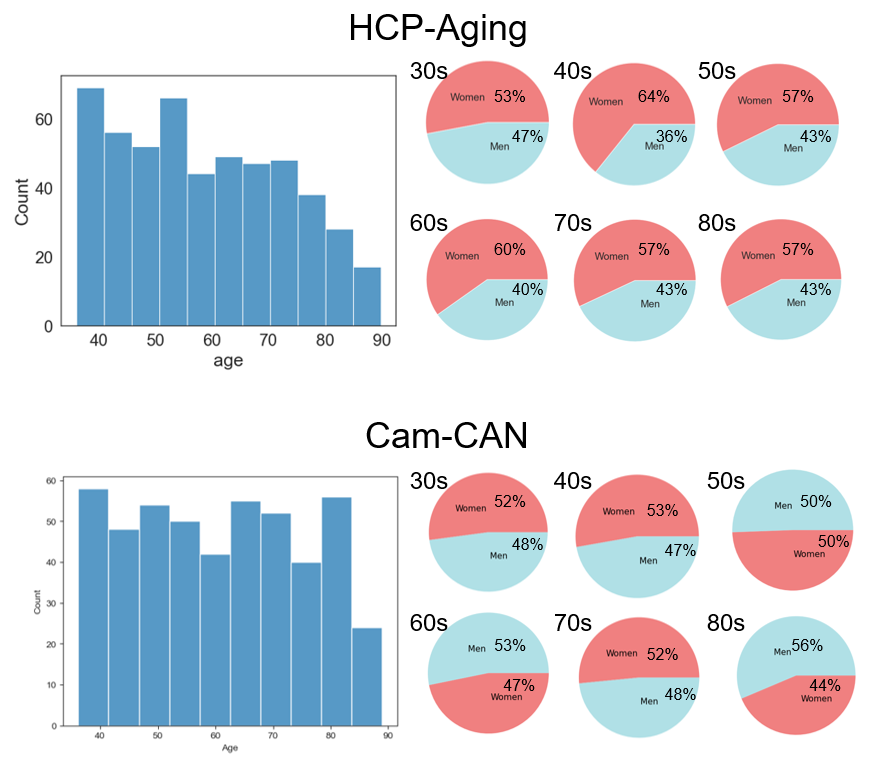

**Figure S1. Age and Gender distribution of HCP-Aging and Cam-CAN dataset.** The figure indicates that in both datasets there is some variance in how many participants are in each decade of life and that it is slightly more women participating in both datasets. Pie charts indicate the distribution of women (red) to men (blue) for each decade, while histograms show the general distribution of ages present in the datasets.

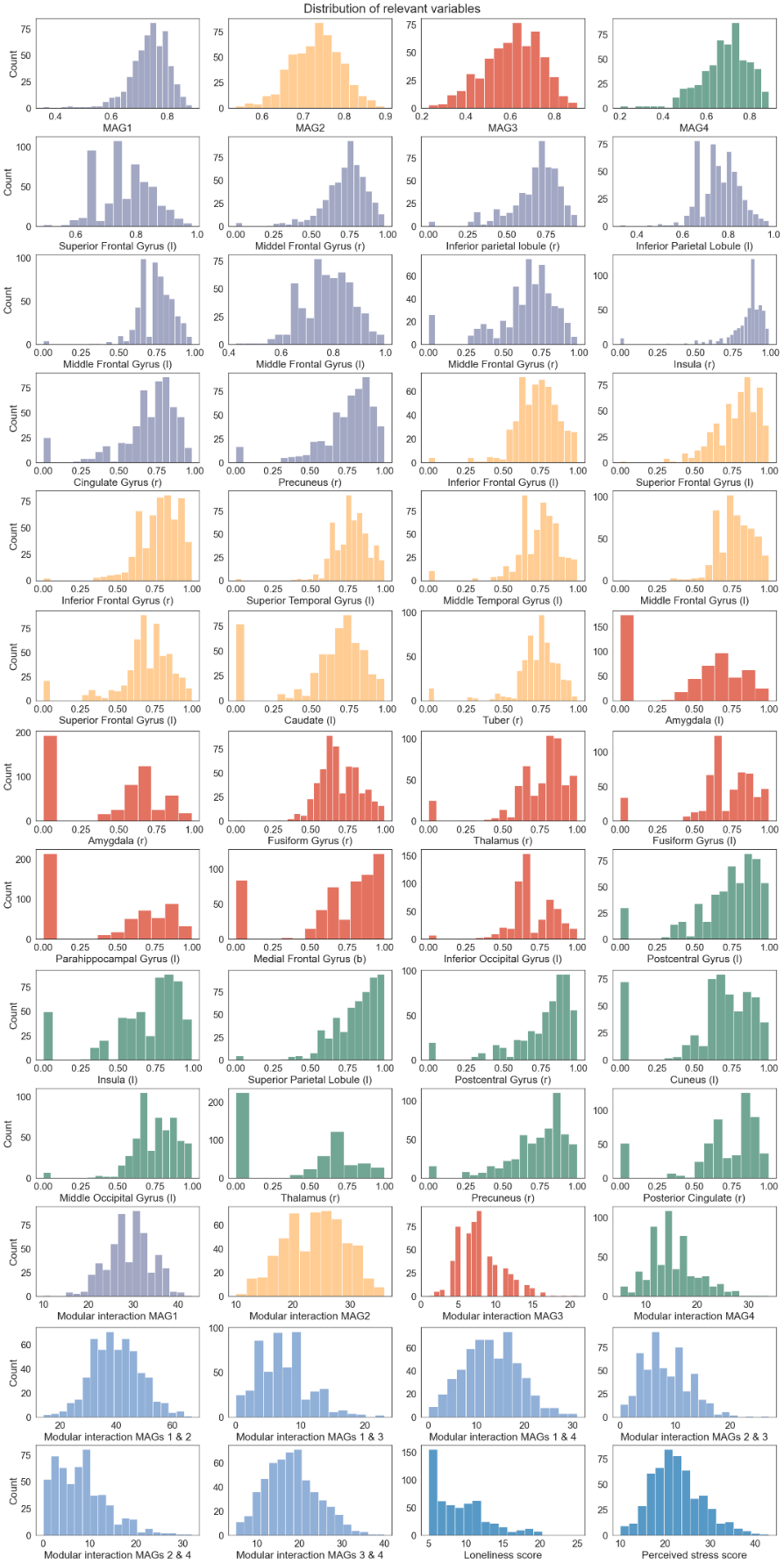

**Figure S2. Histograms of variables.** The distribution of values over every relevant variable used in the analyses. Purple: MAG1 or area belonging MAG1. Yellow: MAG2 or area belonging MAG2. Red: MAG3 or area belonging MAG3. Green: MAG4 or area belonging MAG4. Light blue: interaction between MAGs. Dark blue: behavioral assessment variable. The network PC values seem to follow a normal distribution, while on the nodal level, the distributions are sometimes skewed or the value 0 occurs often.

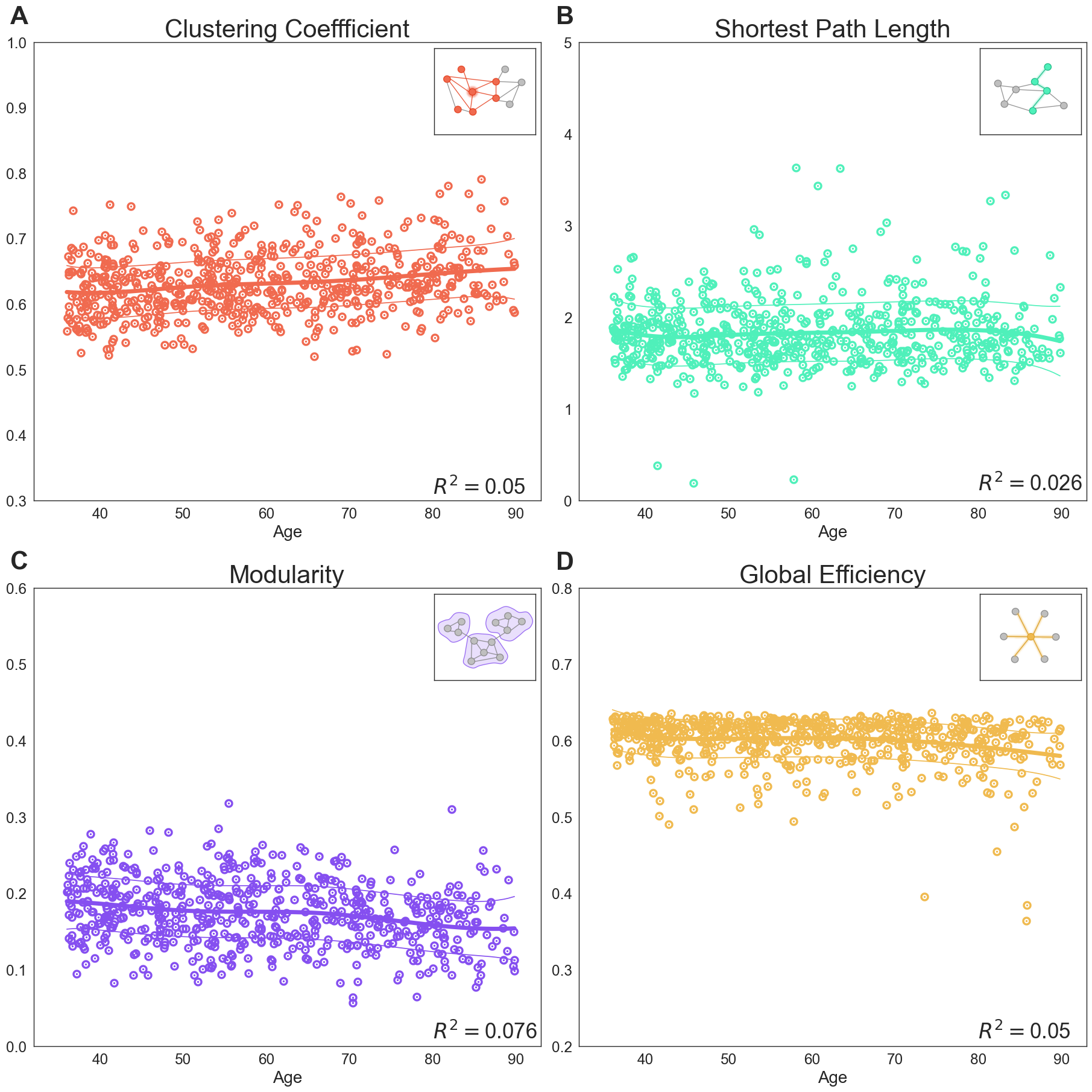

**Figure S3.** GAM fits of additional measurements to assess integration of MAG networks: (A) The clustering coefficient across age shows a mostly linear increase with age, indicating higher integration of the MAGs with one another, while (B) the shortest path length and global network efficiency (D) is mostly unchanged with age. (C) The modularity of the four MAGs shows a significant decrease across the adult life span, further supporting the evidence that large-scale brain networks become more integrated with age. The small boxes symbolize what feature is shown across different ages. R2 = coefficients of determination derived from the general additive models (GAMs).

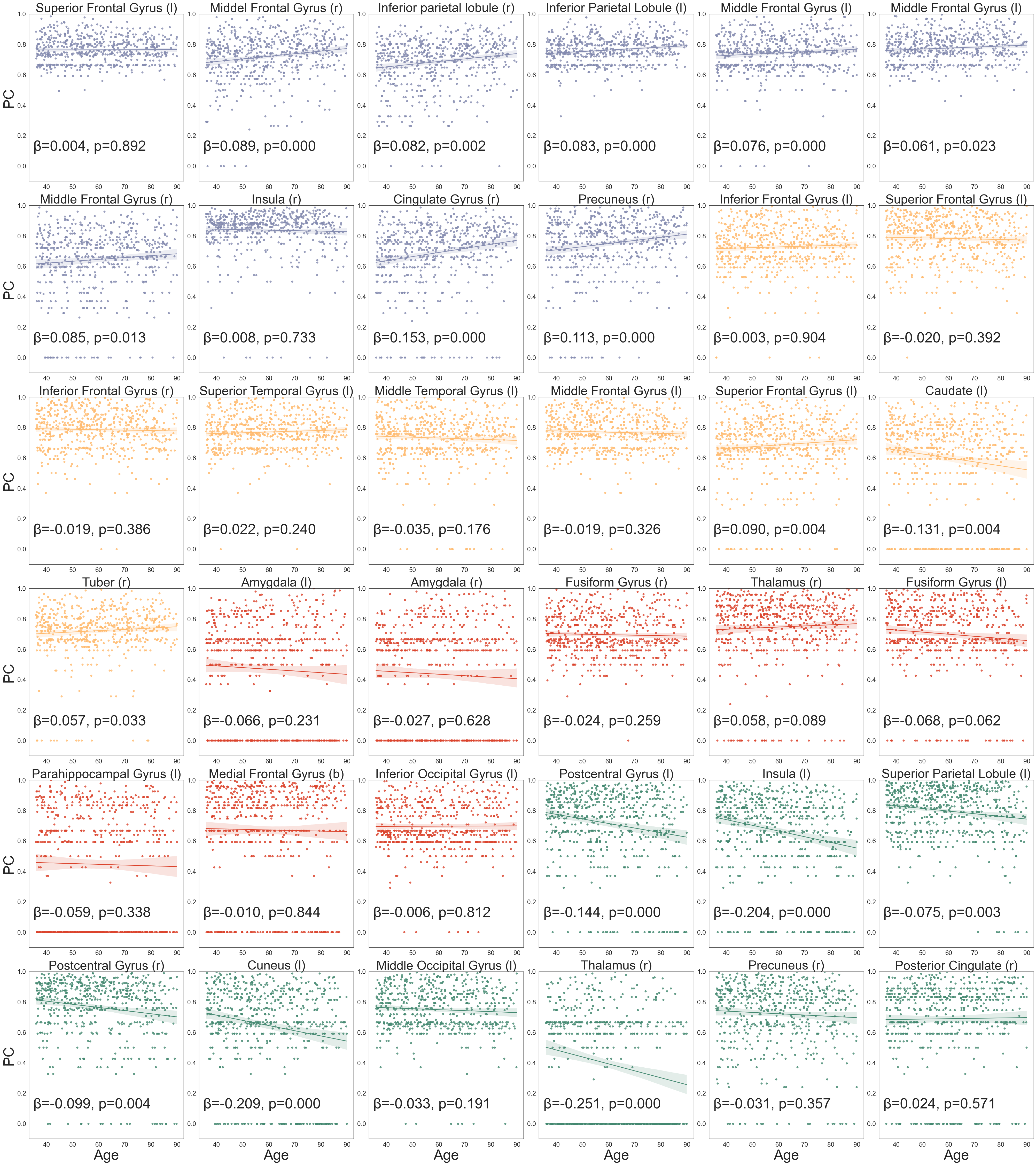

**Figure S4. Linear regression models for the integration of each brain node with age:** purple: areas belong to MAG1, yellow: areas belong to MAG2, red: areas belong to MAG3, green: Areas belong to MAG4. In MAG1 most areas showed increased integration, this increase was significant for the right and left middle frontal gyrus, the right and left inferior parietal lobule, the right cingulate and the right precuneus. The integration of ROIs of MAG2 and MAG3 are mostly stable across different ages, except a decrease in integration of the caudate of MAG2. For MAG4 most areas follow a negative trend in integration with age, a significant decrease was present in the left and right postcentral gyrus, left insula, left superior parietal lobule, left cuneus, and right thalamus.

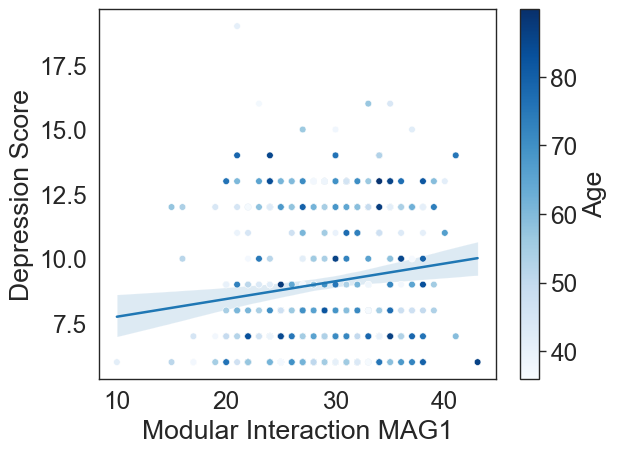

**Figure S5: Association Modular Interaction of MAG1 with Depression Scores.** The inter-connectivity of MAG1 shows a positive association (*r*=0.142, *p*<0.001) with depression score irrespective of age. Each dot is a subject with the darkness of the color representing the subject’s age. Darker color indicates higher age. All ages are distributed evenly across the values. It is important to mention that these are healthy subjects, thus most depression scores remain in the low range.

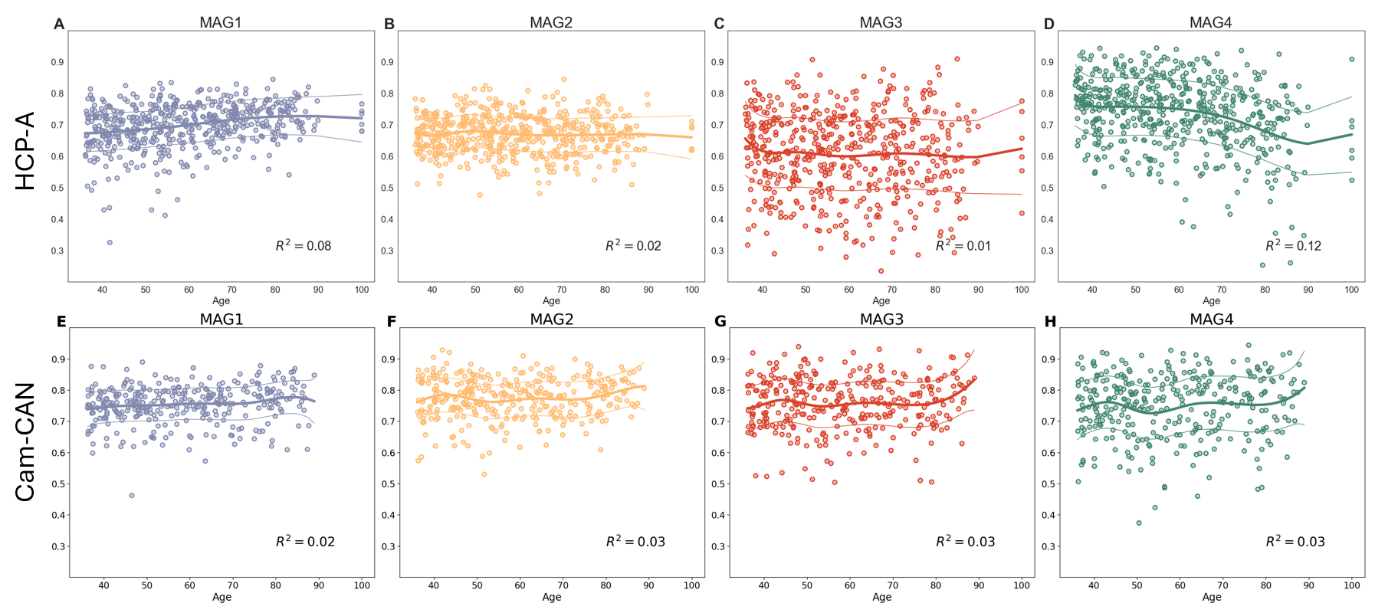

**Figure S6: GAM fit on the network PC of the MAGs in HCP-Aging and Cam-CAN Datasets**. The fitted generalized additive model (GAM) for the integration of the four MAGs. (A-D) GAM fits on the HCP-data. (E-H) GAM fits on the Cam-CAN data which replicated the finding of increased PC of MAG1 in later life (compare plots A & E).
